# Infectious Disease Dynamics Inferred from Genetic Data via Sequential Monte Carlo

**DOI:** 10.1101/096396

**Authors:** R. A. Smith, E. L. Ionides, A. A. King

## Abstract

Genetic sequences from pathogens can provide information about infectious disease dynamics that may supplement or replace information from other epidemiological observations. Currently available methods first estimate phylogenetic trees from sequence data, then estimate a transmission model conditional on these phylogenies. Outside limited classes of models, existing methods are unable to enforce logical consistency between the model of transmission and that underlying the phylogenetic reconstruction. Such conflicts in assumptions can lead to bias in the resulting inferences. Here, we develop a general, statistically efficient, plug-and-play method to jointly estimate both disease transmission and phylogeny using genetic data and, if desired, other epidemiological observations. This method explicitly connects the model of transmission and the model of phylogeny so as to avoid the aforementioned inconsistency. We demonstrate the feasibility of our approach through simulation and apply it to estimate stage-specific infectiousness in a subepidemic of HIV in Detroit, Michigan. In a supplement, we prove that our approach is a valid sequential Monte Carlo algorithm. While we focus on how these methods may be applied to population-level models of infectious disease, their scope is more general. These methods may be applied in other biological systems where one seeks to infer population dynamics from genetic sequences, and they may also find application for evolutionary models with phenotypic rather than genotypic data.

## 1 Introduction

Phylodynamic methods extract information from pathogen genetic sequences and epidemiological data to infer the determinants of infectious disease transmission (Grenfell *et al*., 2004). For successful phylodynamic inference, mechanisms of transmission must leave their signature in genetic sequences. This occurs when pathogen transmission and evolution occur on similar timescales (Drummond *et al*., 2003). By explicitly relating models of disease dynamics to their predictions with respect to pathogen sequences, it is possible to estimate aspects of the mechanisms of transmission (Frost *et al*., 2015; Poon, 2015; Rasmussen *et al*., 2011; Stadler *et al*., 2013; Volz *et al*., 2013b). So far as we are aware, all existing phylodynamic inference methods proceed in three stages. First, one estimates the pathogen phylogeny using sequence data. Next, one fits models of disease dynamics to properties of the pathogen phylogeny, such as coalescent times or summary statistics on the tree. Finally, one assesses the robustness of the results to variation in the estimated phylogeny to account for phylogenetic uncertainty. Frequently, such methods harbor logical inconsistencies between the assumptions of the model used to estimate the phylogeny and those of the model of disease dynamics. In particular, it may happen that population dynamics, as estimated by the transmission model, are inconsistent with those assumed when estimating the phylogeny. In the absence of consistent methods, it may be difficult to assess the loss of accuracy due to the use of inconsistent methods.

Phylodynamic inference for simple deterministic population models is feasible via Markov chain Monte Carlo (Bouckaert *et al*., 2014). However, population models currently computationally feasible via this approach are simple compared to typical models of epidemiological interest (Vaughan *et al*., 2014). A comparable sequential Monte Carlo method applicable to deterministic population models also been proposed (Bouchard-Côté *et al*., 2012). The practice of phylodynamic inference for infectious diseases has therefore focused on methodology conditioning on a phylogeny estimated using a different, simpler model (Frost *et al*., 2015; Karcher *et al*., 2016).

In this paper, we develop methodology for jointly inferring both phylogeny and transmission, as well as estimating unknown model parameters. Our central contribution is an algorithm which we call GenSMC, an abbreviation of *sequential Monte Carlo with genetic sequence data*. Sequential Monte Carlo (SMC), also known as the particle filter, provides a widely used basis for inference on complex dynamic systems (Kantas *et al*., 2015) with several appealing properties. Because basic SMC methods rely only on forward-in-time simulation of stochastic processes, it can accommodate a wide variety of models: essentially any model that can be simulated is formally admissible. Thus, the algorithm enjoys a variant of the plug-and-play property (Bretó *et al*., 2009; He *et al*., 2010). An SMC computation results in an evaluation of the likelihood, which is a well-understood and powerful basis for both frequentist and Bayesian inference. Finally, again because SMC requires only forward-in-time computation, it is straightforward to construct a model of genetic sequence evolution upon the basis of a transmission model, thus avoiding all conflict between these models.

SMC techniques have previously been used for phylodynamic inference conditional on a phylogeny (Rasmussen *et al*., 2011). However, using SMC to solve the joint inference problem through forward-intime simulation of tree-valued processes is a high-dimensional, computationally challenging problem. We found that several innovations were necessary to realize a SMC approach that is computationally feasible on models and datasets of scientific interest. The key innovations that provided a path to feasibility were: just-in-time construction of state variables, hierarchical sampling, algorithm parallelization, restriction to a class of physical molecular clocks, and maximization of the likelihood using the iterated filtering algorithm of Ionides *et al*. (2015).

In the following, we first give an overview of the class of models for which our sequential Monte Carlo algorithms are applicable. A formal specification is given in the supplement, and the source code for our implementation is also available. Next, we present a study on a simulated dataset as evidence of the algorithm’s feasibility. Finally, we use our methods to estimate determinants of the epidemic of human immunodeficiency virus (HIV) among young, black, men who have sex with men (MSM) population in Detroit, Michigan from 2004 to 2011. This analysis uses time-of-diagnosis and consensus protease sequences to estimate the rates of infection attributable to sources inside and outside the focal population.

## 2 New Approaches

The key novelty in our approach to phylodynamics is in formulating a flexible class of phylodynamic models and a class of sequential Monte Carlo algorithms in such a way that the latter can be efficiently applied to the former. We refer to our phylodynamic model class as GenPOMP models, in recognition of the fact that they are partially observed Markov processes (POMPs). As such, a GenPOMP model consists of an unobserved Markov process—called the latent process—and an observable process. In the following sections, we specify the structure of each of these components. An addition, more formal, description of the GenPOMP model is given in the supplement (Section S1). Our GenSMC algorithm for GenPOMP models is introduced in the Materials and Methods section. GenSMC is presented at greater length in the supplement (Section S2) and also provided with a mathematical justification (Section S3). Our extension of GenSMC to parameter estimation, via iterated filtering, is called the GenIF algorithm and is discussed briefly in the Material and Methods section and at greater length in Section S2.2. For computational implementation of the GenPOMP framework and the GenSMC and GenIF algorithms, we wrote the open-source `genPomp` program discussed further in Section S2.1.

For concreteness, we focus here on an infectious disease scenario, wherein the model describes transmission of infections among hosts and the sequences come from pathogens in those infections. In this context, measurements on infected individuals are called diagnoses. In the concluding discussion section, we briefly consider other contexts within which the models and methods we have developed may prove useful.

### 2.1 The latent process

We adopt the convention of denoting random variables using uppercase symbols; we denote specific values assumed by random variables using the corresponding lowercase symbol. We use an asterisk to denote the data, which are treated as a specific realization of random variables in the model.

The latent Markov process, {*X*(*t*), *t* ∈ 𝕋}, defined over a time interval 𝕋 = [*t*_0_, *t*_end_], explicitly models the population dynamics and also includes any other processes needed to describe the evolution of the pathogen. Specifically, we suppose that we can write *X*(*t*) = (𝒯(*t*), 𝒫(*t*), 𝒰(*t*)), where 𝒯(*t*) is the *transmission forest*, 𝒫(*t*) is the *pathogen phylogeny* equipped with a relaxed molecular clock, and 𝒰(*t*) represents the state of the pathogen and host populations. For example, *U*(*t*) may categorize each individual in the host population into classes representing different stages of infection. We suppose that {*U*(*t*),*t* ∈ 𝕋} is itself a Markov process.

The transmission forest represents the history of transmission among hosts. 𝒯(*t*) is a (possibly disconnected) directed, acyclic graph. Nodes in 𝒯(*t*) are time-stamped and of several types. Internal nodes represent transmission events. Terminal nodes are of three types: (a) *active nodes* represent infections active at time *t*; (b) *observed nodes* correspond to diagnosis events, possibly associated with genetic sequences; (c) *dead nodes* correspond to death or emigration events. Root nodes at time *t*_0_ correspond to infections present in the initial population; root nodes at times *t* > *t*_0_ correspond to immigration events. Since all nodes are time-stamped, edges of 𝒯(*t*) have lengths measured in units of calendar time.

The pathogen phylogeny 𝒫(*t*) represents the history of divergences of pathogen lineages. Internal nodes of 𝒫(*t*) represent branch-points of pathogen lineages, which, we assume, coincide with transmission events. The terminal nodes of 𝒫(*t*) are in 1-1 correspondence with the terminal nodes of 𝒯(*t*). The distinction between 𝒫(*t*) and 𝒯(*t*) allows for random variation in the rate of molecular evolution, i.e., relaxed molecular clocks (see below). Specifically, the edge lengths of 𝒯(*t*) measure calendar time between events, whereas edge lengths in 𝒫(*t*) can have additional random variation describing non-constant rates of evolution.

The transmission forest 𝒯(*t*) can grow in only five distinct ways: (1) active nodes can split in two, when a transmission event occurs, (2) active nodes can become dead nodes, upon emigration, recovery, or death of the corresponding host, (3) immigration events can give rise to new active nodes, each with its own distinct root, (4) sampling events cause active nodes to spawn diagnosis nodes, and (5) active nodes for which none of the above occur simply grow older. Likewise, the pathogen phylogeny 𝒫(*t*) grows along with 𝒯(*t*) (Fig. 1). The Markov process {𝒰(*t*)} can contain additional information about the system at time t, e.g., states of individual hosts. {𝒰(*t*)} can affect, but must not be affected by, the {𝒯(*t*)} and {𝒫(*t*)} processes. That is, given any sequence of times *t*_1_ < … < *t_k_* < *t*, {𝒰(*t*)} is independent of {(𝒯(*t_j_*), 𝒫(*t_j_*)), *j* =1,…,*k*} conditional on {𝒰(*t_j_*), *t*_1_ < … < *t_k_* < *t*}.

**Figure 1:**
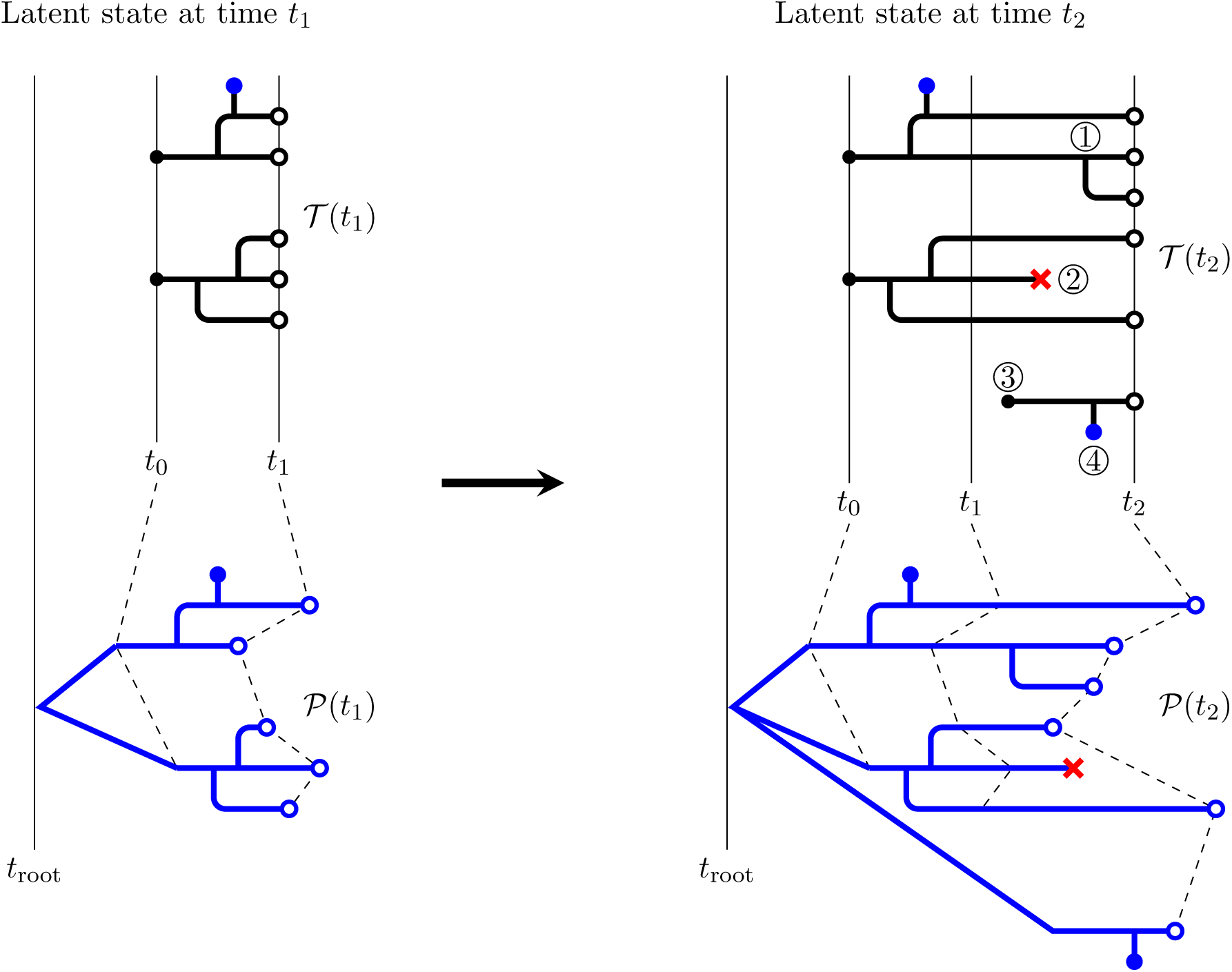
A schematic showing the nature and evolution of the latent transmission and phylogeny processes. The transmission forest, 𝒯(*t*), is shown in black; the pathogen phylogeny, 𝒫(*t*), in blue. At left, we see the latent state at time *t*_1_; it evolves by time *t*_2_ to the state shown at right. At time *t*_1_, 𝒯(*t*_1_) consists of two disconnected trees, representing the transmission histories of five active infections (∘). These infections derive from two infections present at *t*_0_ (black dots). The branching pattern of the pathogen phylogeny mirrors that of 𝒯(*t*) over the interval [*t*_0_, *t*_1_]. This diagram assumes that pathogen lineages branch exactly at transmission events; alternative models could allow for differences in the branching pattern between 𝒯(*t*) and 𝒫 (*t*). This diagram displays a model with a relaxed molecular clock; randomness in the rate of evolution along lineages is depicted via random edge lengths in 𝒫(*t*). Over the time interval [*t*_1_, *t*_2_], changes of each of the five permissible types are shown. At ①, an active node splits in two when a transmission event occurs. At ②, an active node becomes a dead node (×) when an infected host emigrates, recovers, or dies. At ③, an immigration event gives rise to a new active node with its own root. At ④, a sequence node (●) is spawned when a sample is taken. Finally, active nodes for which none of the above occur simply persist. The Markovian property insists that the latent state at time *t*_2_ be an extension of the latent state at time *t*_1_. In other words, changes to the latent state over the interval [*t*_1_,*t*_2_] must not retroactively modify elements of the latent state prior to time *t*_1_.

We assume subsequently that 𝒫(*t*) and 𝒯(*t*) agree topologically, but we note that this assumption is not essential. In particular, the sequential Monte Carlo algorithms we apply could be straightforwardly extended to allow the topology and timing of genetic lineage divergences to deviate from those of transmission events and to allow multiple pathogen lineages within each host. Such extensions might be useful, for example, in accounting for within-host pathogen diversity.

### 2.2 The observable process

We now describe the model explicitly linking the latent process to the data. Let 𝕐 be the set of all finite collections of dated genetic sequences, with an element of 𝕐 being a collection {(*g_k_*, *t_k_*), *k* = 1,…, *n*} where *g_k_* is a sequence and *t_k_* is the associated diagnosis time. We allow *g_k_* to be an empty sequence, in the event that the corresponding diagnosis had no associated sequence. The observable process is a 𝕐-valued process, {*Y*(*t*), *t* ∈ 𝕋}, where *Y*(*t*) consists of all sequences that have accumulated up to time *t*. Thus, *Y*(*t*) is expanding, i.e., *Y*(*t*) ⊂ *Y*(*t′*) whenever *t* ≤ *t′*, and if *Y*(*t*) = {(*G_k_*, *T_k_*), *k* = 1,…, *N*}, then *T_k_* ≤ *t* for all *k*. The data are modeled as a realization of the observable process, *Y*(*t*_end_) = *y*\*.

Suppose each diagnosis has has an equal and independent chance to give rise to a pathogen sequence, and each diagnosis event in *Y*(*t*) corresponds to a unique diagnosis node in 𝒯(*t*). Suppose also that some time-reversible molecular substitution model is defined to describe sequence evolution on the pathogen phylogeny 𝒫(*t*). These modeling assumptions implicitly define a conditional distribution for *Y*(*t*) given *X* (*t*).

### 2.3 Relaxed molecular clocks

A strict molecular clock assumes that the rate of evolution is constant through time and across lineages. Relaxation of this assumption has been shown to improve the fit of phylogenetic models to observed genetic sequences in many cases (Drummond *et al*., 2006) and for HIV in particular (Posada and Crandall, 2001). A relaxed molecular clock models the rate of evolution as random. In our approach, this corresponds to constructing each edge length of 𝒫(*t*) as a stochastic process on the corresponding edge of 𝒯(*t*). Various forms of such processes have been assumed in the literature (Ho and Duchne, 2014; Lepage *et al*., 2007), but not all of these are compatible with a mechanistic approach. In particular, a mechanistic molecular clock must be defined at all times and must have non-negative increments. Many relaxed clocks commonly employed in the literature do not enjoy the latter property: in effect, such clocks allow evolutionary time to run backward. The class of suitable random processes includes the class of nondecreasing Lóevy processes, i.e., continuous-time processes with independent, stationary, non-negative increments.

### 2.4 The plug and play property

The formulation of the latent and observable processes as above is flexible enough to embrace a wide range of individual-based models. In particular, models that describe actual or hypothetical mechanisms of transmission and disease progression are readily formulated in this framework. Moreover, with this formulation, it becomes clear that the models described are partially observed Markov processes (Bretó *et al*., 2009). This fact makes sequential Monte Carlo methods for likelihood-based inference available for use in the present context. The supplementary material makes the formal connections between this class of models and sequential Monte Carlo methodology.

It is worth noting that models formulated as above are compatible with inference techniques that only require simulation from the model, not closed-form expressions for transition probabilities. Such algorithms are said to have the *plug-and-play* property (Bretó *et al*., 2009; He *et al*., 2010). The particle filter and iterated filtering, which we describe in the Methods section, are two algorithms that have this property. Because these algorithms only require the ability to simulate from the model, they allow for consideration of a wide class of models. Greater freedom in choice of the form of the model allows one to pose scientific questions closed to non-plug-and-play approaches. In the following, we demonstrate this potential in a study of HIV transmission dynamics.

## 3 A model of HIV transmission

Our study focuses on the expanding HIV epidemic among young, black, MSM within the Detroit metropolitan area. Specifically, we ask two questions: (1) How much transmission originates inside the study population relative to that originating outside? (2) Within the study population, how does transmission vary with respect to stage of disease (e.g., early, chronic, AIDS) and diagnosis status? To address these questions we construct a basic model of HIV transmission, similar to that of Volz *et al*. (2013a). We describe our model as a special case of the general class of models described above. This model contains assumptions that can be altered and examined within our methodological framework. In the following, we describe both the form of the model and how we relate it to two data types: diagnosis times and genetic sequences.

### 3.1 The latent and observable processes

The latent state of the system at time *t*, (𝒯(*t*), 𝒫(*t*), 𝒰(*t*)), is of the form described above. To specify it completely, it remains to describe the Markov process {𝒰(*t*)} and the transitions of {𝒯(*t*)} and {𝒫(*t*)}. 𝒰(*t*) contains information about all infected individuals in the population. Following Volz *et al*. (2013a), we do not explicitly track uninfected individuals and thus disallow depletion of the susceptible pool. Specifically, 𝒰(*t*) = {(τ_*i*_,*B_i_*(*t*)) : *i* infected at time *t*}, where τ_*i*_ is the time at which individual *i* was infected and *B_i_*(*t*) ∈ ℂ, the class of individual *i* at time *t*, where ℂ = {*I*_0_, *I*_1_, *I*_2_, *J*_0_, *J*_1_, *J*_2_}. *B_i_*(*t*) = *I_k_* indicates that individual *i* has an infection at stage *k* ∈ {0,1,2} but has not yet been diagnosed; *B_i_*(*t*) = *J_k_* indicates that individual *i* has been diagnosed and has an infection at stage *k*. We think of *k* = 0 as indicating the early stage of infection; *k* = 1, the chronic stage; *k* = 2, AIDS. Individuals move between classes according to Fig. 2. New infections can occur, as can deaths, emigrations, and diagnosis events. Transmission events, immigration events, deaths, and diagnoses all result in events of the corresponding type being recorded in the structure of 𝒯(*t*).

**Figure 2:**
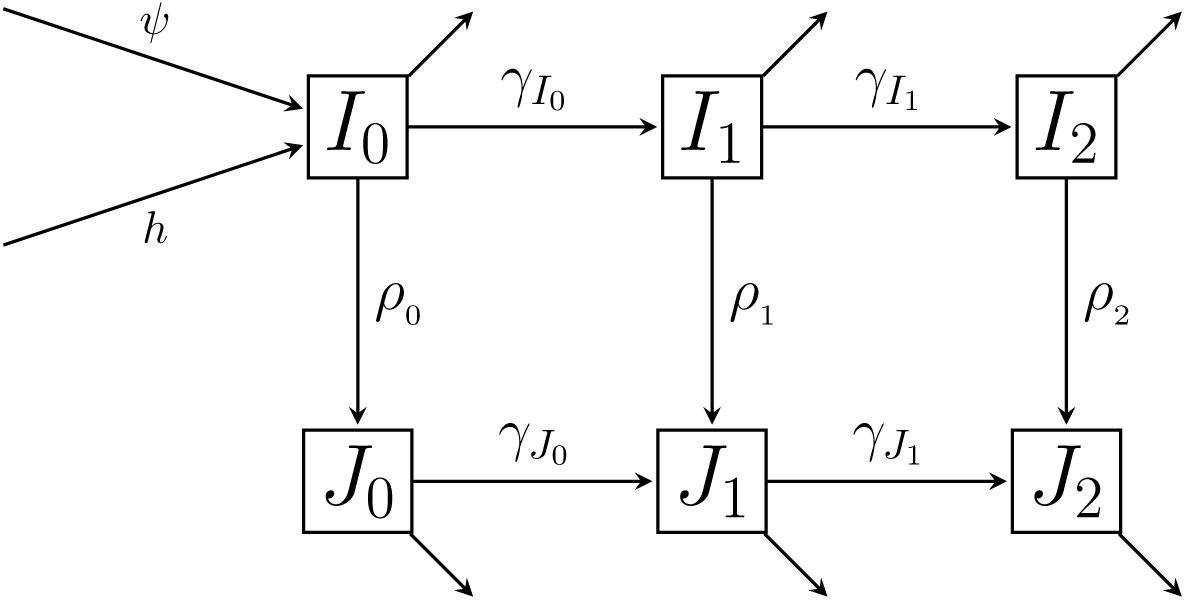
A flow diagram showing the possible classes for infected individuals. The columns represent stage of disease: with subscripts 0, 1 and 2 representing early, chronic, and AIDS stages respectively. The rows represent diagnosis status, with the top row representing undiagnosed individuals, *I_k_*, and the bottom row representing diagnosed individuals, *J_k_*, where *k* ∈ {0,1, 2}. *ρ_k_* are per capita rates of diagnosis and γ_*c*_ are rates of disease progression. Arrows out of classes that do not flow into other classes represent the combined flow out of the infected population due to death and emigration.

New infections arise from two distinct sources: immigration and transmission within the population. Immigrations occur at a constant rate, *ψ*. Each currently infected individual inside the population seeds new infections at rate *ε*_0_, where *c* ∈ ℂ indicates infection class. Thus, we allow transmissibility to vary between different infection classes, but assume homogeneous transmissibility within each class. It follows that the incidence of new infections is *h*(*t*) + *ψ*, where *h*(*t*) = *ε*_*I*_0__ *N*_*I*_0__ (*t*) + *ε*_*I*_1__ *N*_*I*_1__ (*t*) + *ε* _*I*_2__ *N*_*I*_2__ (*t*) + *ε*_*J*_0__ *N*_*J*_0__ (*t*) + *ε*_*J*_1__ *N*_*J*_1__ (*t*) + *ε* _*J*_2__*N*_*J*_2__ (*t*), and *N_c_*(*t*) is the number of individuals in class *c* at time *t*. Defining all nonzero transition rates between states is sufficient to specify a Markov process; a full set of model equations for {𝒰(*t*)} is presented in the supplement (Section S4).

The inclusion of individual time-of-infection, τ_*i*_, within {𝒰(*t*)} allows us to model within-host pathogen evolution. In particular, when an individual is diagnosed at time *t*, a diagnosis node is added to 𝒯(*t*), together with a diagnosis edge, the length of which is linearly related to how long the diagnosed individual has been infected (Fig. 1). This edge may account for sequencing error; it can also describe the emergence of new pathogen strains within a host having reduced between-host transmission potential (Lythgoe and Fraser, 2012).

We assume for simplicity that the topology of 𝒫(*t*) matches that of 𝒯(*t*). Thus, we explicitly disallow the possibility of incomplete lineage sorting, though, as mentioned before, this choice is not forced by the algorithm. We assume a relaxed molecular clock: the edge lengths of 𝒫(*t*) are random. Specifically, each edge of 𝒫(*t*) has length conditionally Gamma distributed with expectation equal, and variance proportional, to the corresponding edge of 𝒯(*t*). That is, if L is the length of an edge of 𝒫(*t*) corresponding to an edge of length *D* in 𝒯(*t*), we posit that *L*|*D* is Gamma distributed with 𝔼 [*L*|*D* = *d*] = *d* and Var [*L*|*D* = *d*] = *σ d*. The parameter *σ* scales the noise on the rate of evolution. This molecular clock relaxation maintains additivity in evolutionary edge lengths and is the same as the white noise model of Lepage *et al*. (2007). Having specified 𝒫(*t*), the joint distribution of observed sequences is determined by the choice of the time-reversible molecular substitution model. Here, we used the HKY model of molecular evolution (Hasegawa *et al*., 1985).

## 4 Results

We present results from both a study on simulated data and an analysis of actual data. The primary goal of the simulation study is to show how our methods can be used to extract information about transmission dynamics from pathogen genetic sequence data within the framework of likelihood-based inference. This study was carried out with 30 sequences of length 100 bases. The goals of the data analysis are to demonstrate the numerical feasibility of our implementation as well as illustrate the role of likelihood-based inference as part of the cycle of data-informed model development for a phylodynamic model. The data analysis was carried out using 100 protease consensus sequences of length 297 bases. Due to the intensive nature of the computations, further developments will be required to handle considerably larger datasets. Some empirical results concerning how our GenSMC implementation scales with number of sequences are given in the supplement (Section S2.3). We discuss applicability to the range of current phylodynamic challenges in the discussion section.

### 4.1 A study on simulated data

Using the individual-based, stochastic model of HIV described above (Fig. 2), we simulated epidemics conditional on observing 30 sequences. We set the length of the simulated sequences to be 100 bases. We set parameters governing the rate of evolution at relatively high values to generate a high proportion of variable sites. As computation scales with the number of variable sites, the computational effort in this simulation study could be comparable to fitting real sequences of greater length. Parameters values and their interpretations are specified in Tables 1 and 2. Algorithmic parameters are specified in Section S4.2. Each simulated epidemic consisted of a transmission forest and a set of pathogen genetic sequences. We randomly selected 5 epidemics to fit. Each dataset consists of two types of data: times of diagnoses and pathogen genetic sequences. A representative simulated transmission forest and its associated pathogen genetic sequences are shown in Fig. 3.

**Figure 3:**
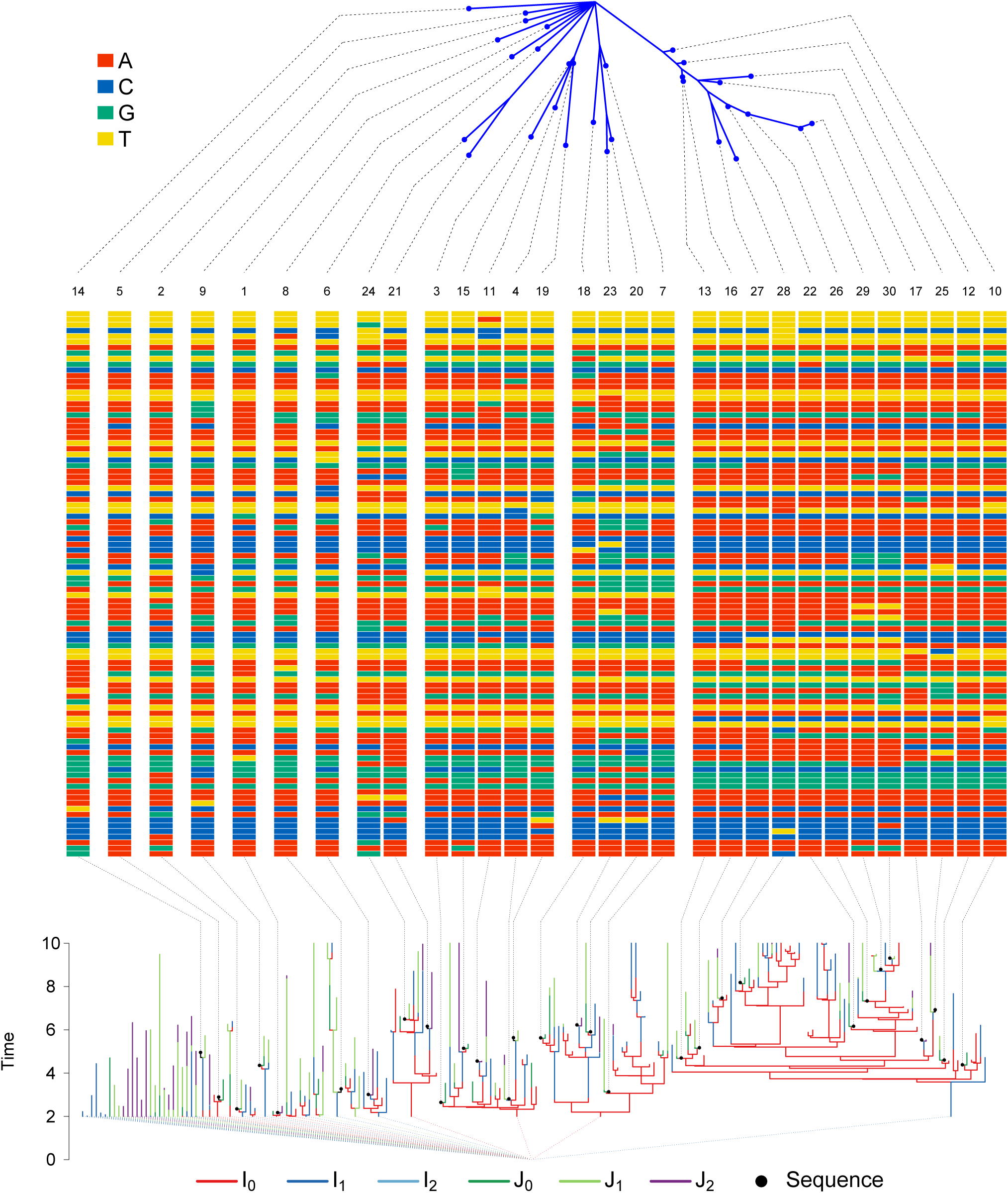
A simulated transmission forest (bottom), its associated pathogen genetic sequences (middle), and the phylogeny of the sequences (top). The class of the infected individual in the transmission forest is indicated by its color. Black dots represent genetic sequences. Black dashed lines connect sequence locations on the transmission tree, or the phylogeny, to visualizations of the sequences in the middle panel. Colored dashed lines from the roots of transmission trees connect at the polytomy at *t_root_* = 0. Numbers at the top of the sequences indicate the rank of the sequence, with rank 1 being the first observed.

**Table 1:** Parameters of the transmission model used in simulation of datasets.

| Parameter | Interpretation | Value |
| --- | --- | --- |
| $\varepsilon_{I_1}$ | Infectiousness of undiagnosed chronic stage individuals | $0.25 \text{ yr}^{-1}$ |
| $\varepsilon_{I_2}$ | Infectiousness of undiagnosed AIDS individuals | $0 \text{ yr}^{-1}$ |
| $\varepsilon_{J_0}$ | Infectiousness of diagnosed acute stage individuals | $0.125 \text{ yr}^{-1}$ |
| $\varepsilon_{J_1}$ | Infectiousness of diagnosed chronic stage individuals | $0.025 \text{ yr}^{-1}$ |
| $\varepsilon_{J_2}$ | Infectiousness of diagnosed AIDS individuals | $0 \text{ yr}^{-1}$ |
| $\mu_{I_0}$ | Death rate + Aging rate of undiagnosed acute stage individuals | $1/3 \text{ yr}^{-1}$ |
| $\mu_{I_1}$ | Death rate + Aging rate of undiagnosed chronic stage individuals | $1/3 \text{ yr}^{-1}$ |
| $\mu_{I_2}$ | Death rate + Aging rate of undiagnosed AIDS individuals | $5/6 \text{ yr}^{-1}$ |
| $\mu_{J_0}$ | Death rate + Aging rate of diagnosed acute stage individuals | $1/3 \text{ yr}^{-1}$ |
| $\mu_{J_1}$ | Death rate + Aging rate of diagnosed chronic stage individuals | $1/3 \text{ yr}^{-1}$ |
| $\mu_{J_2}$ | Death rate + Aging rate of diagnosed AIDS individuals | $2/3 \text{ yr}^{-1}$ |
| $\gamma_{I_0}$ | Progression rate from undiagnosed acute to undiagnosed chronic | $1 \text{ yr}^{-1}$ |
| $\gamma_{I_1}$ | Progression rate from undiagnosed chronic to undiagnosed AIDS | $1/6.3 \text{ yr}^{-1}$ |
| $\gamma_{J_0}$ | Progression rate from diagnosed acute to diagnosed chronic | $1 \text{ yr}^{-1}$ |
| $\gamma_{J_1}$ | Progression rate from diagnosed chronic to diagnosed AIDS | $1/6.3 \text{ yr}^{-1}$ |
| $\rho_0$ | Diagnosis rate of acute stage individuals | $0.5 \text{ yr}^{-1}$ |
| $\rho_1$ | Diagnosis rate of chronic stage individuals | $0.225 \text{ yr}^{-1}$ |
| $\rho_2$ | Diagnosis rate of AIDS individuals | $50 \text{ yr}^{-1}$ |
| $\psi$ | Immigration rate of infected individuals | $0 \text{ yr}^{-1}$ |
| $\phi$ | Emigration rate of infected individuals | $0 \text{ yr}^{-1}$ |
| $t_{\text{root}}$ | Root (polytomy) time | $0 \text{ yr}$ |
| $t_0$ | Time to begin simulation of the transmission model | $2 \text{ yr}$ |
| $t_{\text{end}}$ | Time to end simulation of the transmission model | $10 \text{ yr}$ |
| $n_{\text{loci}}$ | Length of the sequences to simulate | 100 base pairs |
| $p_G$ | Probability of a sequence given diagnosis | 0.48 |
| $N_{I_0}(t_0)$ | Number of undiagnosed early-stage individuals at $t_0$ | 11 |
| $N_{I_1}(t_0)$ | Number of undiagnosed chronic-stage individuals at $t_0$ | 15 |
| $N_{I_2}(t_0)$ | Number of undiagnosed AIDS individuals at $t_0$ | 0 |
| $N_{J_0}(t_0)$ | Number of diagnosed early-stage individuals at $t_0$ | 4 |
| $N_{J_1}(t_0)$ | Number of diagnosed chronic- stage individuals at $t_0$ | 8 |
| $N_{J_2}(t_0)$ | Number of diagnosed AIDS individuals at $t_0$ | 6 |

**Table 2:** Parameters of the genetic model used in simulation of datasets.

| Parameter | Interpretation | Value |
| --- | --- | --- |
| $\beta$ | Rate of transversions | $0.013 \text{ yr}^{-1}$ |
| $\alpha_Y$ | Rate of transitions between purines | $0.03 \text{ yr}^{-1}$ |
| $\alpha_R$ | Rate of transitions between pyrimidines | $0.1 \text{ yr}^{-1}$ |
| $\pi_A$ | Equilibrium frequency of adenine | 0.37 |
| $\pi_G$ | Equilibrium frequency of guanine | 0.23 |
| $\pi_C$ | Equilibrium frequency of cytosine | 0.18 |
| $\pi_T$ | Equilibrium frequency of thymine | 0.22 |
| $\sigma_{\text{site}}$ | Relaxation of the molecular clock with respect to sites | 0 |
| $\sigma$ | Relaxation of the molecular clock with respect to edges | 0.1 yr |
| $\delta_{\text{fixed}}$ | The initial component of the sequence stem | 0.001 yr |
| $\delta_{\text{prop}}$ | Proportion of time since infection to add to the sequence stem | 0.05 |

For each of the selected epidemics we ask two questions. First, when all other parameters are known, is it possible to infer *ε*_*I*_0__ and *ε*_*I*_1__ using only diagnosis times? Second, how does inference change when we supplement the diagnosis data with pathogen genetic sequences? To perform this comparison we estimated two likelihood surfaces for each epidemic: one using only the diagnosis likelihood, and one using both the diagnosis likelihood and the genetic likelihood. We estimated each surface by using the particle filter to compute a grid of likelihood estimates with respect to the two parameters of interest: *ε*_*I*_0__, the infectiousness of early-stage undiagnosed individuals, and *ε*_*I*_1__, the infectiousness of chronic-stage undiagnosed individuals. Equilibrium base frequencies were set to the empirical values in the simulated data. All other parameters were fixed at the known values used for simulation. We extracted grid-based likelihood profiles for each parameter by taking maxima over the columns or rows of the grid. For each parameter we therefore obtained two profiles: one using only the diagnosis likelihood and one using the joint likelihood. The difference in curvature between these profiles tells how much the genetic data improves, or weakens, inference on the parameters.

When only the diagnosis data are used, we find a tradeoff between *ε*_*I*_0__ and *ε*_*I*_1__ (Fig. 4). The diagnoses provide information on upper bounds for each infectiousness parameter, but otherwise only inform their sum. In other words, when estimated using only the diagnosis times, *ε*_*I*_0__ and *ε*_*I*_1__ are nonidentifiable. Supplementing the data on diagnoses with pathogen genetic sequences resolves this uncertainty (Fig. 4). Note that including the genetic data increases noise in the likelihood estimate. This is expected, as computing the likelihood estimate for the genetic sequences requires a numerical approximation to an integral over tree space. Nevertheless, the genetic data increase the curvature of the likelihood surface. From Fig. 4, we see that this additional curvature leads to more precise identification of the parameters despite the increased Monte Carlo noise. In principle, Monte Carlo variation can be reduced to negligibility by increased computational effort. This may not be practical when computational expense is high, as it is here. Therefore, it is necessary to bear in mind the tradeoff between the benefits of the information accessed for inference versus the computational burden of extracting this information.

**Figure 4:**
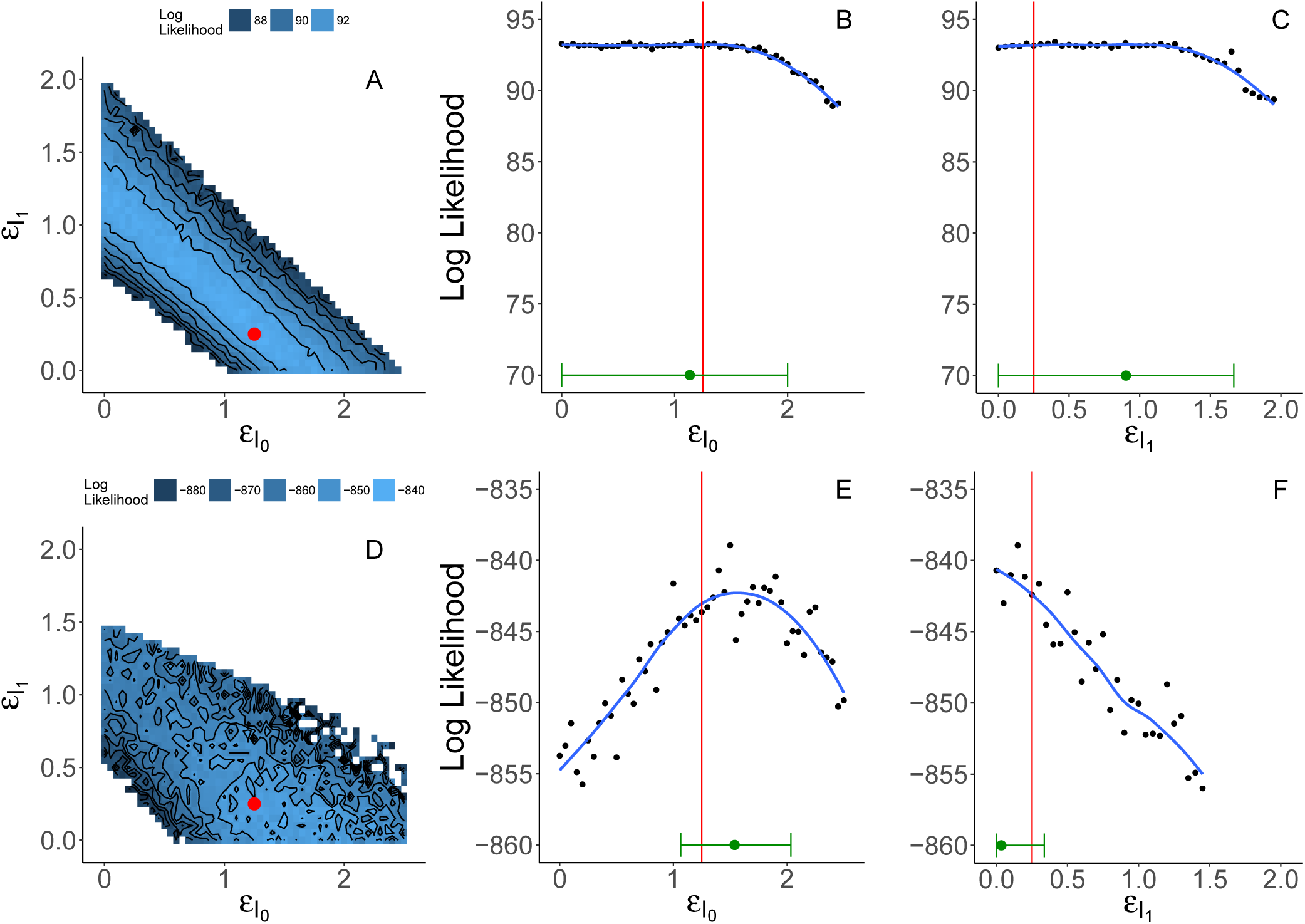
Grid-based estimates of likelihood surfaces and likelihood profiles from fitting to simulated data. The top row shows the surface (A) and profiles (B and C) estimated using only the diagnosis likelihood. The bottom row shows the surface (D) and profiles (E and F) estimated using both the diagnosis and the genetic likelihood. Red dots and red lines indicate true values of *ε*_*I*_0__ and *ε*_*I*_1__ used in simulation. Point estimates and 95% confidence intervals are shown in green just above the horizontal axis of the likelihood profile plots. Confidence intervals for E and F account for both statistical uncertainty and Monte Carlo noise (Ionides *et al*., 2016) using a square root transformation appropriate for non-negative parameters. Augmenting the diagnosis data with genetic data yields smaller confidence intervals for *ε*_*I*_0__ and *ε*_*I*_1__, and resolves the nonidentifiability of these parameters when estimated using only the diagnoses. Note that scales of the likelihood surfaces shown in A and D are not the same; E and F have the same scale as B and C but with a vertical shift.

### 4.2 Analysis of an HIV subepidemic in Detroit, MI

In this data analysis, we explored whether our full-information approach could estimate key transmission parameters using HIV protease consensus sequences and diagnosis times. We focused our analysis on a subepidemic in the young, black, MSM community. The cohort of individuals that we chose to study is shown in Fig. 5. See the Materials and Methods section for details on how we selected the subepidemic and cleaned the sequence data.

**Figure 5:**
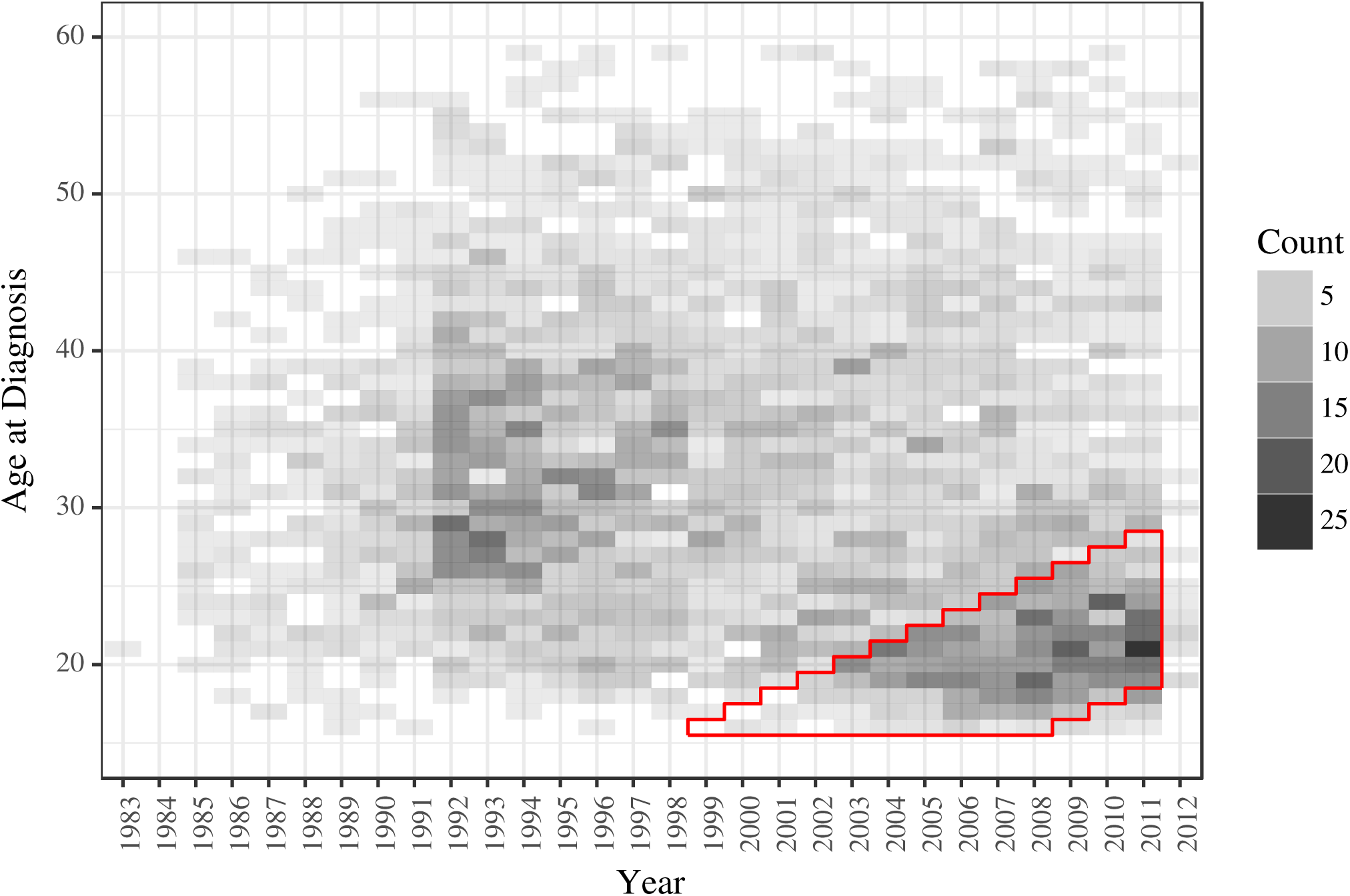
The distribution of age at diagnosis through time for black MSM in Detroit, MI. The cohort that we selected for analysis is outlined in red. We excluded the data from 2012 to limit effects from delays in updating the MDCH database. 29 individuals that were diagnosed at ages greater than or equal to 60 years are not shown on this plot.

As in the study on simulated data, we were interested in what the genetic data yield beyond what we can see using the diagnoses alone. Therefore, we again estimated likelihood profiles in two ways: using only the diagnosis data and using both the diagnosis data and the genetic sequences. We estimated likelihood profiles for three parameters of interest: *ε*_*I*_0__, *ε*_*J*_0__, and *ψ*. In contrast to the simulation study, in this analysis we were faced with a parameter space of much higher dimension. To reduce the dimension of the problem we fixed some parameters: rates of disease progression, rates of diagnosis, and the rate of emigration. Parameters that were fixed and fit are shown in Tables 3 and 4, respectively. Algorithmic parameters are specified in Section S4.2. For each likelihood profile we first used iterated filtering Ionides *et al*. (2015) to maximize the likelihood for a sequence of values that spanned the reasonable range of the parameter. Second, we used the particle filter to estimate likelihoods for each parameter set obtained from iterated filtering. We repeated this process of maximization followed by evaluation until the profile stabilized. All initial-value parameters were fixed, with the exception of *t*_root_. Initial counts for individuals in each class were fixed. See the supplement for details on how we arrived at these counts.

**Table 3:** Parameters fixed in the data analysis.

| Parameter | Interpretation | Value |
| --- | --- | --- |
| $\mu_{I_0}$ | Death rate of undiagnosed acute stage individuals | $1/70 \text{ yr}^{-1}$ |
| $\mu_{I_1}$ | Death rate of undiagnosed chronic stage individuals | $1/70 \text{ yr}^{-1}$ |
| $\mu_{I_2}$ | Death rate of undiagnosed AIDS individuals | $1/2 \text{ yr}^{-1}$ |
| $\mu_{J_0}$ | Death rate of diagnosed acute stage individuals | $1/70 \text{ yr}^{-1}$ |
| $\mu_{J_1}$ | Death rate of diagnosed chronic stage individuals | $1/70 \text{ yr}^{-1}$ |
| $\mu_{J_2}$ | Death rate of diagnosed AIDS individuals | $1/70 \text{ yr}^{-1}$ |
| $\gamma_{I_0}$ | Progression rate from undiagnosed acute to undiagnosed chronic | $1 \text{ yr}^{-1}$ |
| $\gamma_{I_1}$ | Progression rate from undiagnosed chronic to undiagnosed AIDS | $1/6.3 \text{ yr}^{-1}$ |
| $\gamma_{J_0}$ | Progression rate from diagnosed acute to diagnosed chronic | $1 \text{ yr}^{-1}$ |
| $\gamma_{J_1}$ | Progression rate from diagnosed chronic to diagnosed AIDS | $1/6.3 \text{ yr}^{-1}$ |
| $\rho_0$ | Diagnosis rate of acute stage individuals | $0.225 \text{ yr}^{-1}$ |
| $\rho_1$ | Diagnosis rate of chronic stage individuals | $0.225 \text{ yr}^{-1}$ |
| $\rho_2$ | Diagnosis rate of AIDS individuals | $50 \text{ yr}^{-1}$ |
| $\phi$ | Emigration rate of infected individuals | $0 \text{ yr}^{-1}$ |
| $N_{I_0}(t_0)$ | Number of undiagnosed early-stage individuals at $t_0$ | 20 |
| $N_{I_1}(t_0)$ | Number of undiagnosed chronic-stage individuals at $t_0$ | 36 |
| $N_{I_2}(t_0)$ | Number of undiagnosed AIDS individuals at $t_0$ | 0 |
| $N_{J_0}(t_0)$ | Number of diagnosed early-stage individuals at $t_0$ | 4 |
| $N_{J_1}(t_0)$ | Number of diagnosed chronic- stage individuals at $t_0$ | 22 |
| $N_{J_2}(t_0)$ | Number of diagnosed AIDS individuals at $t_0$ | 16 |
| $\sigma_{\text{site}}$ | Relaxation of molecular clock with respect to sites | 0 yr |
| $t_0$ | Time to start filtering | 1 Jan 2004 |

**Table 4:** Parameters fit in the data analysis. We present confidences intervals for parameters for which we computed likelihood profiles. For all other parameters, we present only the point estimate.

| Parameter | Interpretation | Diagnosis data | Diagnosis data and genetic sequences | Diagnosis data, with $\psi$ fixed at 0 |
| --- | --- | --- | --- | --- |
| $\psi$ | Immigration rate of infected individuals | 120 (104, 134) yr<sup>-1</sup> | 5.82 (2.55, 11.2) yr<sup>-1</sup> | 0 yr<sup>-1</sup> |
| $\varepsilon_{I_0}$ | Infectiousness of undiagnosed acute stage individuals | 0 (0, 0.413) | 0.257 (0.0399, 0.623) | 0 (0, 0.192) |
| $\varepsilon_{I_1}$ | Infectiousness of undiagnosed chronic stage individuals | 0.0042 | 0.00048 | 0.0056 |
| $\varepsilon_{I_2}$ | Infectiousness of undiagnosed AIDS individuals | 0 | 0 | 0 |
| $\varepsilon_{J_0}$ | Infectiousness of diagnosed acute stage individuals | 0.0675 (0, 1.17) | 3.36 (3.13, 4.2) | 7.34 (5.78, 9.25) |
| $\varepsilon_{J_1}$ | Infectiousness of diagnosed chronic stage individuals | 0.0089 | 0.17 | 0.032 |
| $\varepsilon_{J_2}$ | Infectiousness of diagnosed AIDS individuals | 0 | 0 | 0 |
| $\beta$ | Rate of transversions | - | 0.0042 yr<sup>-1</sup> | - |
| $\alpha_Y$ | Rate of transitions between purines | - | 0.047 yr<sup>-1</sup> | - |
| $\alpha_R$ | Rate of transitions between pyrimidines | - | 0.043 yr<sup>-1</sup> | - |
| $\pi_A$ | Equilibrium frequency of adenine | - | 0.37 | - |
| $\pi_G$ | Equilibrium frequency of guanine | - | 0.24 | - |
| $\pi_C$ | Equilibrium frequency of cytosine | - | 0.18 | - |
| $\pi_T$ | Equilibrium frequency of thymine | - | 0.21 | - |
| $\sigma$ | Relaxation of molecular clock with respect to edges | - | 2 yr | - |
| $\delta_{\text{prop}}$ | Proportion of time since infection to use for diagnosis edge | - | 0.064 | - |
| $\delta_{\text{fixed}}$ | Amount of calendar time to add on to diagnosis edge | - | 0.00049 yr | - |
| $t_{\text{root}}$ | Time of the polytomy that joins all genetic lineages | - | 27 Aug 2000 | - |

When only the diagnosis data are used, we find that the model prefers to explain all infections as originating outside the cohort, with the maximum likelihood estimate (MLE) for *ψ* ≈ 120 infections per year (Fig. 6). Under this explanation for the data, little or no transmission occurs inside the cohort: this covariate-defined subgroup acts as a sentinel of the broader epidemic. Equivalently, this result would imply that the covariates we used to select these cases do not define a meaningful subepidemic.

**Figure 6:**
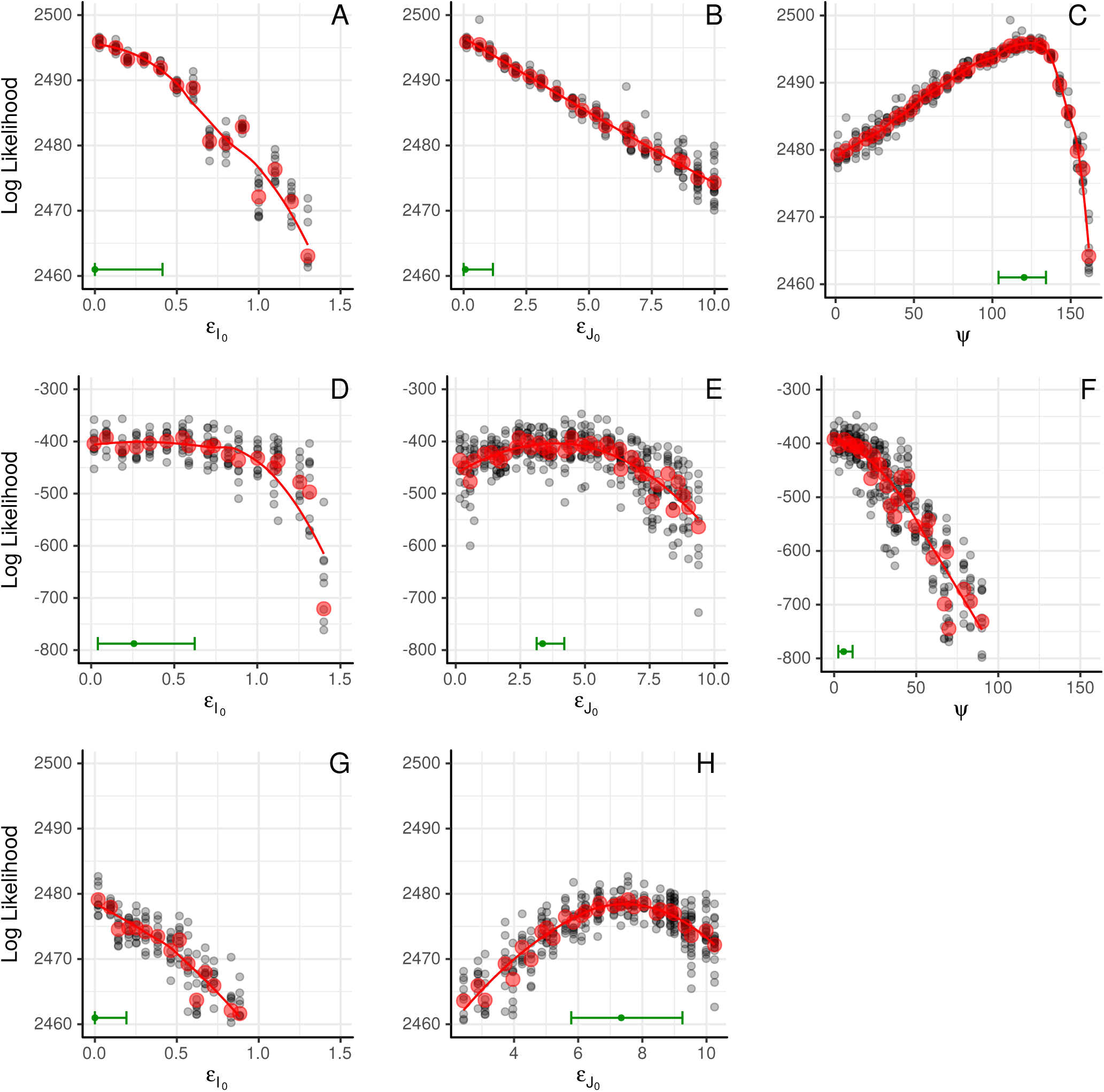
Estimated likelihood profiles from fits to data from the black, MSM cohort. A-C show likelihood profiles computed using only the diagnosis likelihood. D-F show likelihood profiles computed using both the diagnosis likelihood and the genetic likelihood. G and H show likelihood profiles computed using only the diagnosis likelihood when *ψ* is fixed at zero. Black dots represent particle filter likelihood evaluations of parameter sets obtained using iterated filtering. Red dots represent mean log likelihoods of the multiple likelihood evaluations (black dots) at each point in the profile. Red lines are loess fits to the red dots. Green bars along the lower margin of each panel encompass 95% confidence intervals for each parameter. Confidence intervals account for both statistical uncertainty and Monte Carlo noise (Ionides *et al*., 2016). The smoothed profile was calculated on the square root scale, appropriate for non-negative parameters, with a green dot indicating the maximum.

On the other hand, when the genetic data are folded in, the estimate of *ψ* is greatly revised: the MLE for *ψ* becomes ≈ 6 infections per year. On its face, this is evidence for a low rate of transmission into the cohort and, therefore, evidence that the cohort subepidemic is much more self-contained. Although this may in part be true, the lower estimate of *ψ* is also potentially driven by assumptions of the genetic model. Supposing, as it does, that all immigrant lineages coalesce at a single, global polytomy, the model insists that sequences from immigrant infections derive from a broad genetic pool. The breadth of this pool—the average genetic distance between an imported infection and any other observed sequence— is determined by the depth of the polytomy, an estimated parameter. Nevertheless, the low estimate of *ψ* implies that few infections derive from this broader pool. The model’s disallowance of a more structured immigrant pool makes it difficult to say more, however. In particular, the low value of *ψ* is not inconsistent with the existence of chains of transmission originating within the cohort, leaving it, and returning. Such chains would produce sequence clustering despite the openness of the cohort to transmission. Future work, incorporating genetic and diagnosis information from the broader epidemic will be needed to better quantify the latter effect.

Joint likelihood profiles over *ε*_*I*_0__ and *ε*_*J*_0__ show support for transmission from both of the early-stage groups, with evidence for higher infectiousness in the early-stage diagnosed class than in the early-stage undiagnosed class. However, it is epidemiologically implausible that diagnosis increases transmission: this is a paradox. Since the paradox did not arise in the simulation study, it cannot be due to a coding error in the implementation of the model or the statistical methodology. Assuming no errors in the data, therefore, it must derive from some inappropriate feature of the model. We propose two possible explanations for how the model and data combine to yield this result.

One possibility is that temporal clusters of genetically related diagnoses favor high infectiousness for the early-stage diagnosed. For example, this could be an artifact of unmodeled clusters in HIV testing. We searched the data for such clusters, but found no conclusive evidence for their presence.

A second possibility is understood by noting that, under the model, any significant amount of transmission from the undiagnosed classes leads necessarily to an exponentially growing accumulation of diagnoses, in conflict with the data. When the genetic data were left out, the model accounted for the observed, roughly linear, ramp-up in diagnoses using immigration, hence the relatively high estimated *ψ*. Incorporating the genetic data eliminates this option, forcing the model to explain the epidemic’s sub-exponential growth as a consequence of diagnosis itself.

To illustrate the second possibility, we estimated likelihood profiles using only the diagnosis likelihood, fixing the immigration rate, *ψ*, at zero. These profiles show that, when forced to explain the diagnoses without any imported infection, the model prefers to do so by making the early-stage diagnosed class most infectious (Fig. 6). This suggests that the model lacks flexibility to explain the pattern in the diagnoses without immigration; this constraint likely limits efficient use of information in the genetic sequences. To remedy this problem, one could modify the model by explicitly introducing a small and ephemeral population of susceptible hosts.

In this methodological paper, we display but one iteration of the scientific method and it is clear that our motivating scientific questions remain incompletely answered. Our principal goal, however, is to illustrate how the methodology facilitates the formulation and testing of scientific hypotheses. For example, the results above suggest a number of straightforward model modifications: the plug-and-play property of the methodology makes it nearly as straightforward to evaluate the evidence for these new hypotheses just as we have done for the old. Moreover, we have shown how probing the data with a mechanistic model can lead to clear identification of flaws in model structure, along with indications for improvements.

## 5 Discussion

We demonstrated, via a simulation study, that our algorithms provide access to the likelihood surface of a population dynamic model fit to genetic sequence data. This opens the door to likelihood-based phylodynamic inference. As this study shows, incorporating information from genetic data has the potential to improve on inference that we obtain using diagnosis data alone.

In our analysis of an HIV subepidemic in Detroit, MI, we showed that our methods can be used to ask questions of current public health interest by fitting practical models to data of nontrivial size. This study illustrates how the ability to confront the model with different data types, alone or in combination, can be essential to understanding how the model interacts with the data, to uncovering shortcomings of the model, and to pointing the way toward improved model formulations. The ability of our methods to incorporate different data types made it possible to assess each source of information’s contribution to the overall inference. In turn, the ability to easily restructure the model, guaranteed by the plug-and-play property, will allow us to push forward model development.

The scope of our methodology goes beyond the examples presented: the algorithms described here are applicable across a wide range of host-pathogen systems and may find application in realms beyond genetics. From an abstract perspective, these algorithms provide the ability to relate demographic processes with a growing tree-like structure to the evolution of discrete characters that are carried and passed along the branches of that tree. So long as this evolution occurs on a similar timescale to that of the demographic process, and measurements of the discrete process are heterochronous, the methods presented here apply.

In this paper, we demonstrated the methods using relatively short consensus sequences derived from Sanger sequencing. While our methods may be well suited to analysis of data from fast-evolving RNA viruses, they may also apply in studies of pathogens that evolve more slowly. Advances in sequencing are increasing the range of problems for which phylodynamic inference is applicable (Biek *et al*., 2015). The ability to apply phylodynamic inference to bacterial and protozoan genomes opens the door to many epidemiological applications. One area that may be particularly interesting to explore using our methods is hospital outbreaks of drug resistant bacteria. Hospital records on location and duration of stay may provide fine-scale information on populations of susceptible and infected individuals. Accurate measures of these demographic quantities may allow for efficient use of information held in genetic data. Furthermore, the relatively small size of outbreaks in hospitals means that stochasticity may play a large role in their dynamics, and our methods are designed to explicitly account for the role of different sources of stochasticity.

We conclude by placing our new methodology in the context of the eight current challenges identified by Frost *et al*. (2015) for inferring disease dynamics from pathogen sequences. We will make some relevant comments on each challenge, in order.

1. **Accounting for sequence sampling patterns.** Our methodology explicitly models sequence sampling. The chance of an individual being diagnosed, or subsequently having their pathogen sequenced, is permitted to depend on the state of the individual. This state could contain geographic information, or whatever other aspect of the sampling procedure one desires to investigate. Sampling issues revolve around how the dynamics and the measurement process affect the relatedness of sequences, and are more naturally handled in a framework that deals jointly with estimation of the population dynamics and the phylogeny. Thus, our main innovation of joint estimation is directly relevant to this challenge.
2. **Using more realistic evolutionary models to improve phylodynamic inferences.** In this paper, we have used simple evolutionary models that have been widely used for previous phylodynamic inference investigations. Our methodology does not particularly facilitate the use of more complex evolutionary models, since the large number of trees under consideration puts a premium on rapid likelihood computation. However, our methodology is primarily targeted at drawing inference on the population dynamics rather than the micro-evolutionary processes. For this purpose, it may be sufficient to employ an evolutionary model which captures the statistical relationship between genetic distance and temporal distance on the transmission tree, together with an appropriate estimate of the uncertainty in this relationship. Better evolutionary models would be able to extract information more efficiently from the data, but from our perspective this challenge may not be a primary concern.
3. **The role of stochastic effects in phylodynamics.** Our methodology explicitly allows for stochastic effects in the population dynamics and sequence collection.
4. **Relating the structure of the host population to pathogen genetic variation.** Our framework explicitly models this joint relationship. Further scientific investigations, fitting models using methods accounting properly for the joint relationship, will lead to progress in understanding which aspects of dynamics (such as super-spreading) might be especially important to include when carrying out phylodynamic inference.
5. **Incorporating recombination and reassortment.** In principle, our methodology is flexible enough to include co-infection and its evolutionary consequences. Due to computational considerations, it will be important to capture parsimoniously the key aspects of these processes.
6. **Including phenotypic as well as genotypic information.** Our framework naturally combines genotypic information with other information sources. For example, in our data analysis we complemented genetic sequence data with diagnosis times for unsequenced patients.
7. **Capturing pathogen evolution at both within-host and between-host scales.** The diagnosis edges on our phylogenetic tree allow for differences between observed and transmissible strains, and therefore give a representation of within-host diversity or measurement noise. Other approaches to within-host pathogen diversity are possible within our general framework. For example, one could include within-host branching of the phylogenetic tree. More complete investigation of within-host pathogen dynamics will require additional modeling. Due to the larger models and datasets involved, applying our methodology to such investigations will require further methodological work on scaling.
8. **Scaling analytical approaches to keep up with advances in sequencing.** In this manuscript, our goal was to develop generally applicable and statistically efficient methodology. Our methodology is structured with computational efficiency in mind, subject to that goal. Our approach combines various algorithms that have favorable computational properties: peeling, particle filtering with hierarchical resampling and just-in-time variable construction, and iterated filtering. There is scope for computational enhancement by adapting the methodology to high performance architectures. In particular, parallel particle filtering is an active research topic (Paige *et al*., 2014) that is directly applicable to our methodology. There are also possibilities for improving scaling by imposing suitable situation-specific approximations; for example, it might be appropriate to reduce the computational burden by supposing that some deep branches in the phylogeny are known.

In summary, our new methodology has potential for making progress on many of the challenges identified by Frost *et al*. (2015). Beyond that, the methodology offers a full-information, plug-and-play approach to phylodynamic inference that gives the scientist flexibility in selecting appropriate models for the research question and dataset at hand. Although technical challenges remain, especially in scaling these methods to large data, these algorithms hold the potential to ask and answer questions not accessible by alternative approaches.

## 6 Materials and methods

### 6.1 Overview of sequential Monte Carlo estimation of the likelihood

Sequential Monte Carlo (SMC) is a stochastic algorithm originally designed to estimate imperfectly observed states of a system via a collection of dynamically interacting simulations Arulampalam *et al*. (2002). SMC is also called the particle filter algorithm, and each simulation is usually called a particle. SMC sequentially estimates the unobserved state of the system at the time of each observation by iterating through three steps: (1) for each particle, simulate the latent process forward in time to the next data point, (2) for each particle, compute the conditional probability density of the observation given the proposed latent state, and (3) resample the particles with replacement with probabilities proportional to their conditional probabilities. While inference of unobserved states is one use of the particle filter, we are primarily interested in using the filter as an estimator of the likelihood. The average of the conditional likelihoods across particles is an estimator of the conditional likelihood of each observation, and the product of these conditional likelihoods is an unbiased estimator of the full likelihood of the data (Theorem 7.4.2 on page 239 in Del Moral (2004)).

The basic particle filter described above only requires the ability to simulate realizations of the latent state and the ability to evaluate the density of an observation given the latent state. As we explained earlier, in our case the latent state contains both the full transmission forest and the phylogeny of the pathogen lineages. At minimum, the observations consist of a set of genetic sequences of the pathogen. Although in principle these methods could be applied to homochronous sequences, we primarily envision using them to fit models to heterochronous sequences. Additional datatypes can be incorporated into the likelihood evaluation if desired so long as there is a means to relate these data to the latent state.

We implemented the particle filter such that the algorithmic code is independent of the code that specifies the model. This structure allows for realizing the advantanges of the plug-and-play paradigm by facilitating quick comparisons between models of different forms. Pseudocode for the algorithm is provided in the supplement. In Alg. 1 we give an outline of the pseudocode, and we show a schematic of simplest form of the algorithm in Fig. 7. In our framework, the user specifies the model by writing three functions:

1. **A simulator for the initial state of the latent process.** This function initializes 𝒯(*t*_0_). For example, in a model with only one class of infected individuals, this function would initialize 𝒯(*t*_0_) by specifying the number of infected individuals at *t*_0_. Each of these individuals then becomes a root of a tree in the transmission forest. Each root of the transmission forest has its own genetic lineage; these comprise 𝒫(*t*_0_). In our implementation, the initializer does not construct 𝒫 (*t*_0_); the structure of 𝒫(*t*) is built as needed (see below in the section ‘Just-in-time construction of state variables’).
2. **A forward simulator for the latent state.** This function simulates growth of 𝒯(*t*) forward in time from one observation to the next. This function also places the next observation on 𝒯(*t*), assigning the sequence to an individual by augmenting 𝒯(*t*) with a diagnosis edge and a sequence node. Note that this function does not simulate evolution of genetic sequences. Rather, the algorithm proposes ancestral relationships between genetic sequences via the simulated transmission forest. While formally, the pathogen phylogeny 𝒫(*t*) is part of the latent state, for computational efficiency we choose not to simulate its structure in full. The function in (3) builds the necessary components of 𝒫(*t*) given the simulated transmission forest and placement of sequences on the forest.
3. **An evaluator for the conditional probability of observing a sequence.** This function returns the conditional probability of observing a sequence given the latent state and all previously observed sequences. In particular, this function conditions on the structure of the subtree of 𝒫(*t*) that connects the observed sequences. The simplest choice for this function is to (1) make the strong assumption that 𝒫(*t*) maps directly onto 𝒯(*t*), and therefore build the phylogeny based strictly on the topology of 𝒯(*t*) and (2) evaluate the conditional likelihood of the genetic sequence using the peeling algorithm (Felsenstein, 1981). These two choices are equivalent to assuming a strict molecular clock. However, one may choose more complicated functions, such as mappings that allow for discrepency between 𝒯(*t*) and 𝒫(*t*) or a relaxed molecular clock, to better match the mechanistic processes that generate real data. The branching pattern of the transmission forest and of the phylogeny may differ for a number of reasons Romero-Severson *et al*. (2014), so there may be strong arguments for allowing for discrepency between these trees.

**Figure 7:**
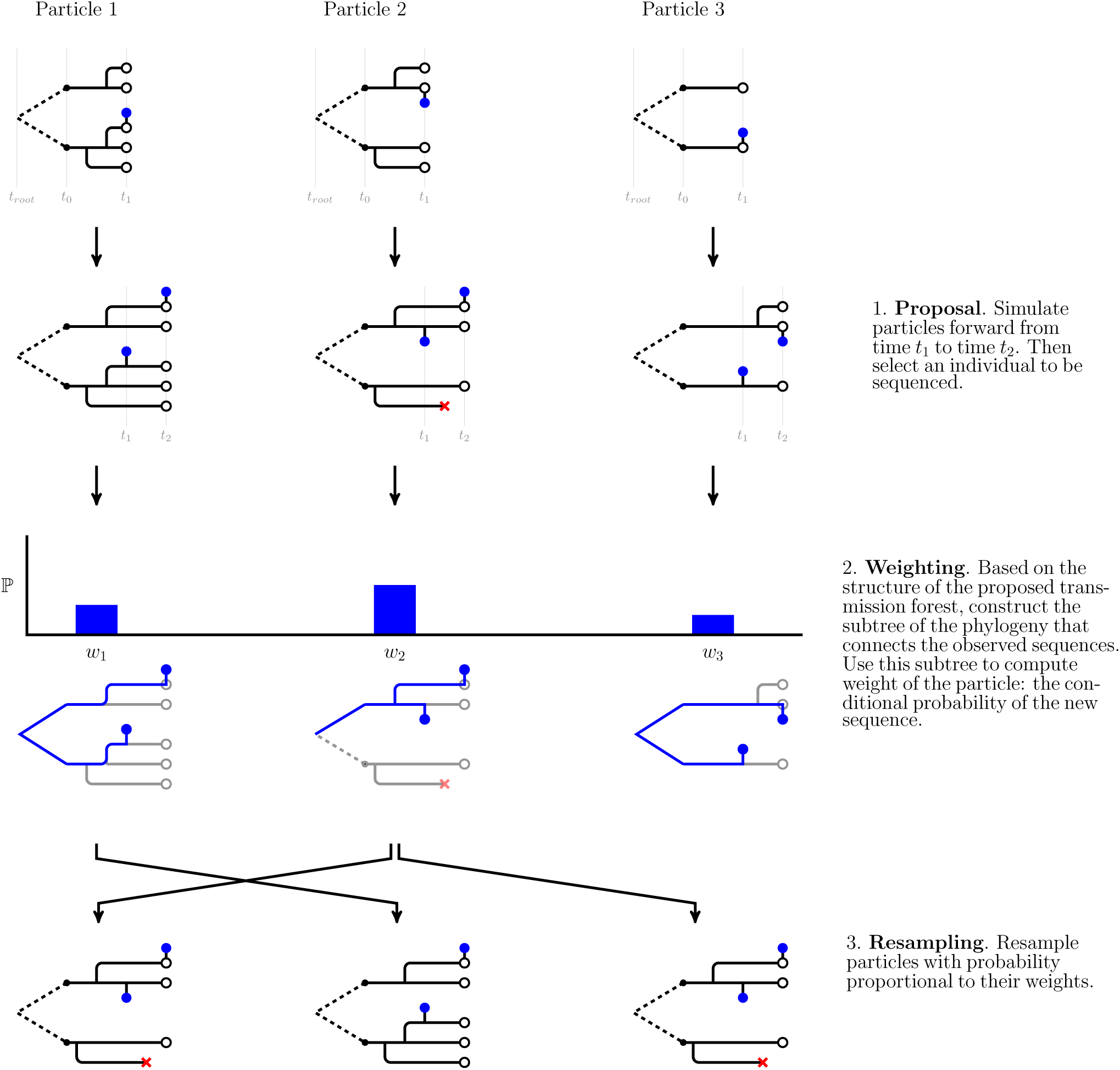
A schematic of the particle filter. Here, we show steps to run the filter from the first sequence to the second. Transmission forests are shown in black and phylogenies that connect observed sequences, 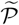(*t*), are shown in blue. Observed sequences are depicted as blue dots. This schematic shows how the algorithm uses *just-in-time* construction of state variables to ease computational costs. Although the model describes how 𝒫(*t*) relates to 𝒯(*t*) across all branches of the transmission tree, the algorithm only constructs the subtree of the phylogeny needed to connect the observations (and therefore evaluate conditional probabilities of sequences). Note that in our implementation of the particle filter we introduce additional procedures in the proposal and weighting steps. These procedures, which are detailed below, allow for more accurate assessment of a particle’s weight (through hierarchical sampling) and estimation of the conditional probability of a sequence under a relaxed clock.

##### Algorithm 1: GenSMC [Corresponding step numbers for the complete description in Section S2 are in brackets]

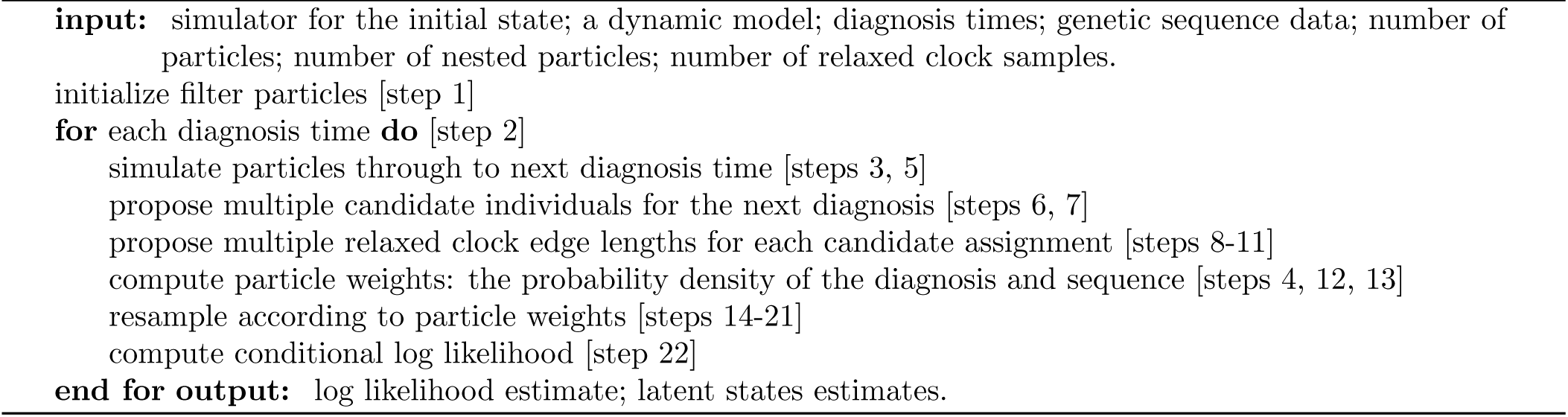

### 6.2 Maximization of the likelihood via iterated filtering

The particle filter provides access to the likelihood surface, but it does not provide an efficient way to maximize the likelihood. A closely related class of algorithms, iterated filtering, allows for maximizing the likelihood. Iterated filtering incorporates perturbation of unknown parameters into the particle filter. Repeatedly passing the filter over the data while shrinking the size of the perturbations allows the parameters to converge to their maximum likelihood estimates. The setup here, with a growing state space, does not perfectly match the framework used to develop iterated filtering by Ionides *et al*. (2015). However, the basic iterated filtering approach of perturbing parameters and filtering repeatedly can be applied, and can be assessed on its empirical success at maximizing the likelihood.

### 6.3 Computational Structure

One way our algorithms differ from a standard SMC approach is that each particle maintains a latent state comprising of tree structures that reach back to *t*_root_. As the algorithm incorporates each additional data point its memory requirement grows. From a practical perspective, the necessity of maintaining a deep structure in the particles presents challenges for writing a computationally feasible implementation of the algorithm. We developed several innovations to meet the computational challenges posed by numerically integrating over tree space. In this section, we give an overview of key components of our implementation that contributed to numerical tractability. For details, see the source code at https://github.com/kingaa/genpomp (to be archived at https://datadryad.org).

#### 6.3.1 Data structures and their relationship to model specification

Our implementation holds two tree structures in memory for each particle: (1) 𝒯(*t*), the transmission tree, and (2) 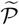(*t*), the subtree of 𝒫(*t*) that connects all sequences observed up to time *t*. We represent 𝒯(*t*) as a vector of nodes, where each node contains the index of its mother, a timestamp, and the index of the genetic lineage with which it is associated (if any). Although the model of the latent state includes the full phylogeny of the pathogen, 𝒫(*t*), our algorithms only need to keep a subtree of the phylogeny, 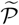(*t*), in memory. We also represent 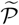(*t*) as a vector of nodes. However, nodes of 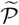(*t*) require more memory than the nodes of 𝒯(*t*). In addition to the information in a transmission tree node, each node of 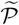(*t*) contains the indices of the node’s daughters, an array of probabilities, and an evolutionary edge length. These additional components allow for computing the likelihood of observing the sequences at the tips of 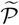(*t*).

Our implementation provides a set of functions that allow for specifying the model via forward-intime simulation of the latent state. These functions provide access to the latent state and allow for modifying the latent state by branching lineages in 𝒯(*t*), terminating leaves in 𝒯(*t*), etc. Our code does not provide access to 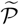(*t*). Instead, internal functions update the structure of 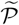(*t*) as necessary (detailed in the following section on just-in-time construction of state variables). The structure of 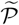(*t*) is in part determined by the molecular clock model. Our current implemention supports strict molecular clock models and relaxed molecular clocks with gamma distributed edge lengths (as we use in this paper). Alternative models for 𝒫(*t*) are possible, and the plug-and-play structure of our algorithms allows the user to explore a wide range of alternative models.

#### 6.3.2 Just-in-time construction of state variables

Although the model of the latent process includes the full phylogeny of the pathogen, 𝒫(*t*), for the purposes of computation we need only store 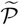(*t*) in memory. In our implementation, we add new edges to 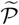(*t*) at the time of measurement; it is not until a sequence is placed on a lineage of 𝒯(*t*) that we have enough information to update 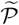(*t*). We call this approach *just-in-time* construction of state variables because simulation of part of the state is postponed until the last moment. An alternative approach would include simulation of 𝒫(*t*) in tandem with the transmission forest. Then, when a sequence is attached to 𝒯(*t*) the necessary components of 𝒫(*t*) to relate the new sequence to all previously observed sequences would be guaranteed to be present. When the transmission forest is large relative to the phylogeny such an approach would be costly in both computation and memory.

#### 6.3.3 A hierarchical sampling scheme

We developed a hierarchical sampling scheme to allow for scaling the effective number of particles while holding only a fraction of the effective number of particles in memory. This sampling scheme allows for holding *J* particles in memory while approaching effective sample sizes approaching *JK*, where *J* is the number of base particles and *K* is the number of nested particles. In this hierarchical scheme, we split the proposal into two steps: (1) proposal of the transmission forest, and (2) proposal of the location of the sampled sequence on the transmission forest. Each of *J* particles first proposes a transmission forest. Then each of the *J* particles calculates the likelihood of the observed sequence for *K* possible locations of the observed sequence (Fig. 8). One of the *K* nested particles is kept, sampled with weight proportional to its conditional likelihood, and the remaining *K* — 1 particles are discarded. The weight of the surviving particle is the average of the conditional likelihoods of the *K* nested particles.

**Figure 8:**
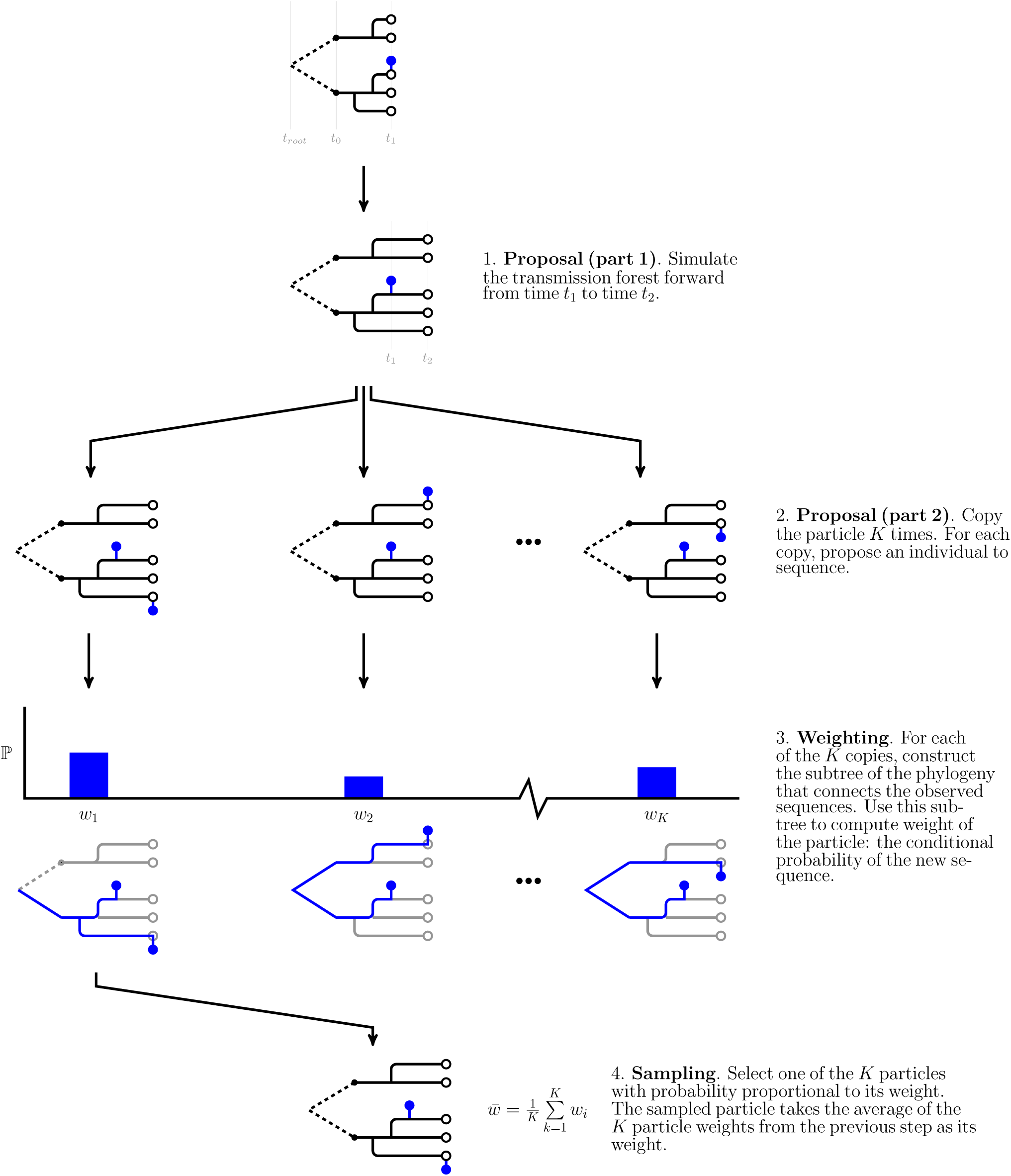
A schematic of our hierarchical sampling scheme. In this scheme, we split the proposal into two steps: (1) simulation of the transmission forest and (2) selecting an eligible individual to be sequenced. When each particle is expensive, it may pay to invest more effort in evaluating the conditional probability of a sequence given the latent state. This procedure is easily nested within the simpler form of the particle filter shown in Fig. 7. In turn, one can add additional Monte Carlo steps to the weighting step in this procedure to evaluate the conditional probability of a sequence under a relaxed clock (see Fig. 9).

#### 6.3.4 A Monte Carlo procedure for the relaxed molecular clock

As we have no closed-form expression for the conditional probability of an observed sequence under a relaxed clock, we estimate this probability via simulation. Fig. 9 shows how we incorporate this Monte Carlo procedure into our SMC framework. We generate *L* instances of the subtree of the phylogeny that connects all previously observed sequences up to time *t*, 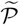(*t*). We then augment each subtree with an edge to accommodate the new sequence. The length of this edge is gamma distributed as described above. When connecting the new edge to the existing phylogeny, there are two cases: either the edge connects at the root or the new edge splits an existing edge. In the case of a split edge, we allocate edge length to either side of the split according to a beta distribution. This procedure maintains gamma distributed edge lengths. Having constructed the phylogeny connecting all sequences up to the new sequence, we then use the peeling algorithm (Felsenstein, 1981) to compute the conditional probability of the new sequence. The average of the conditional probability given each of the *L* subtrees is an estimate of the conditional probability of the new sequence under a relaxed clock.

**Figure 9:**
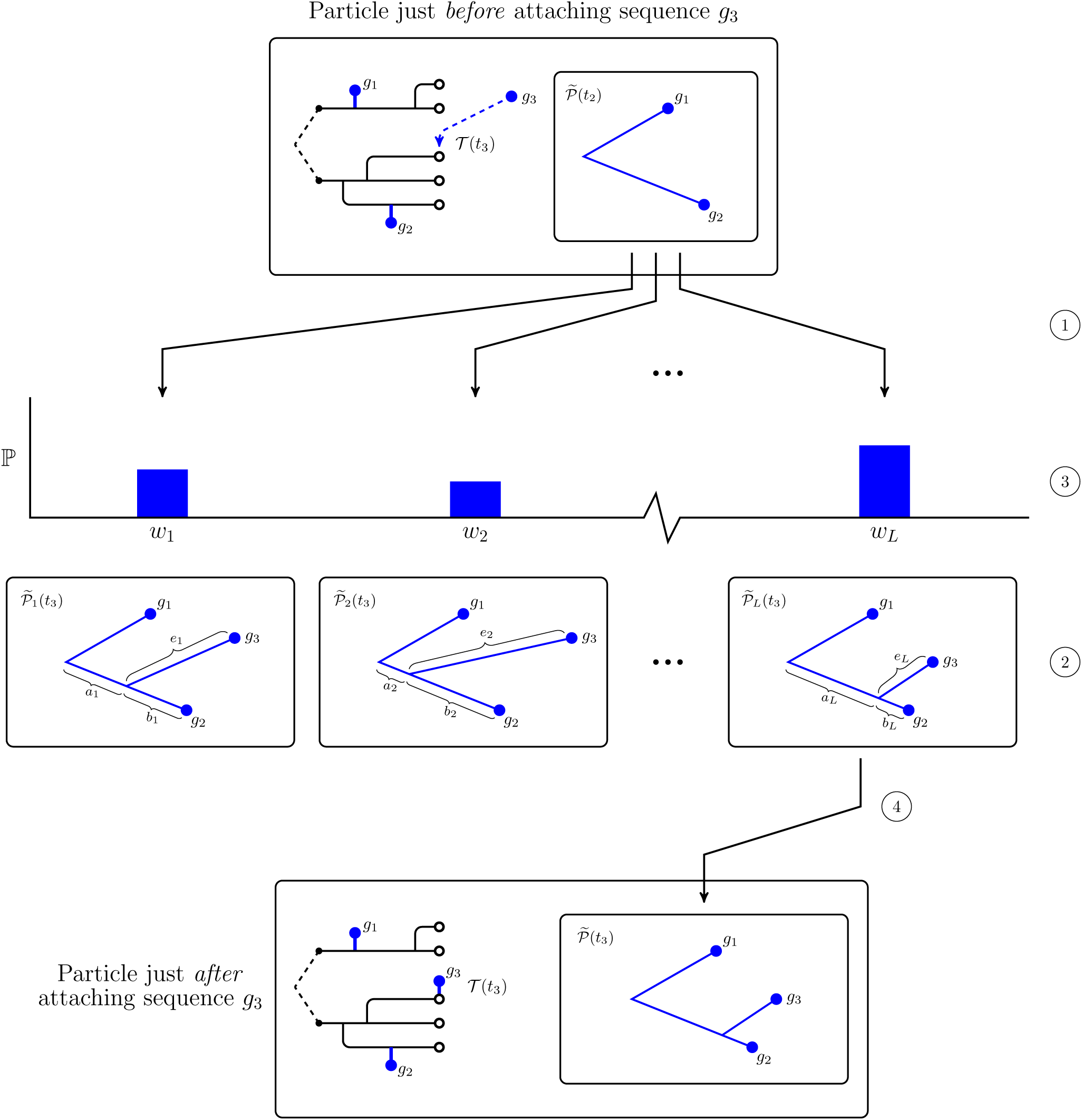
A schematic showing our Monte Carlo approach to estimate the conditional probability of a sequence under a relaxed clock. Note that this procedure only modifies the subtree of the phylogeny that joins the sequences, 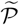(*t*). At the top, we show a particle just before attaching a new sequence. In this case, the particle has already incorporated two sequences, and the location of the third sequence on the transmission forest has already been selected. First, we make *L* copies of 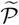(*t*_2_), the subtree of the phylogeny that connects all sequences observed up to time *t*_2_ (at ①). For each of these phylogenies we propose an attachment site and an edge length for sequence *g*_3_ (at ②). The edge length of the edge subtending sequence *g*_3_, *e_l_*, is drawn from a Gamma distribution parameterized as described in the text. We split the edge between the root and sequence *g*_2_ according to a Beta distribution into two lengths, *a_l_* and *b_l_*; this procedure preserves Gamma distributed edge lengths for two components of the split edge. Then, for each proposed phylogeny, we use the peeling algorithm to compute the conditional probability of sequence *g*_3_ (at ③). Finally, we sample one of these proposed phylogenies with probability proportional to its weight (at ④). The unsampled proposals are discarded and the particle takes the average of the conditional probabilities as its weight.

#### 6.3.5 Parallelization

We used openMP (Dagum and Menon, 1998) to parallelize the algorithm at the level of a single machine to reduce runtimes. In particular, we parallelized the outer loop of the hierarchical sampling scheme described above. Each processor handles one base particle at a time. The cost in memory for *n* processors handling *J* particles with a nested sample size of *K* is therefore at worst *J* + *nK*, as each processor may have at most *K* additional particles in memory.

### 6.4 A model of HIV transmission: computation of the measurement model

Each diagnosis event consists of a diagnosis time and, possibly, an associated genetic sequence. In the case where the diagnosis event has no sequence, the measurement model is only the conditional density of the diagnosis time. When there is an associated sequence, it is the product of the conditional density of the diagnosis time and the conditional probability of the genetic sequence.

We compute the conditional density of a diagnosis time as follows. We decompose the density into two terms: (1) The probability of no diagnosis over the last interdiagnosis interval: 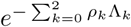 where *ρ_k_* is the diagnosis rate for class *I_k_*, *k* ∈ {0,1, 2}, and Λ_*k*_ is the integrated hazard of a diagnosis from class *I_k_* : Λ_*k*_ = 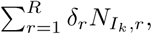 where, *δ_r_* is length of the *r*^th^ subinterval in the interdiagnosis interval over which the count of class *I_k_*, *N*_*I_k_*,*r*_, is constant, and (2) the hazard of a diagnosis at the time of diagnosis: 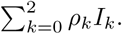 The conditional density of a diagnosis time is the product of these two quantities, and is therefore a mixture of a probability and a density. To compute the first, each particle accumulates the person-years of undiagnosed individuals over the last diagnosis interval (Fig. 10). The second is easily computed given the number of each class of undiagnosed individual at the time of diagnosis.

**Figure 10:**
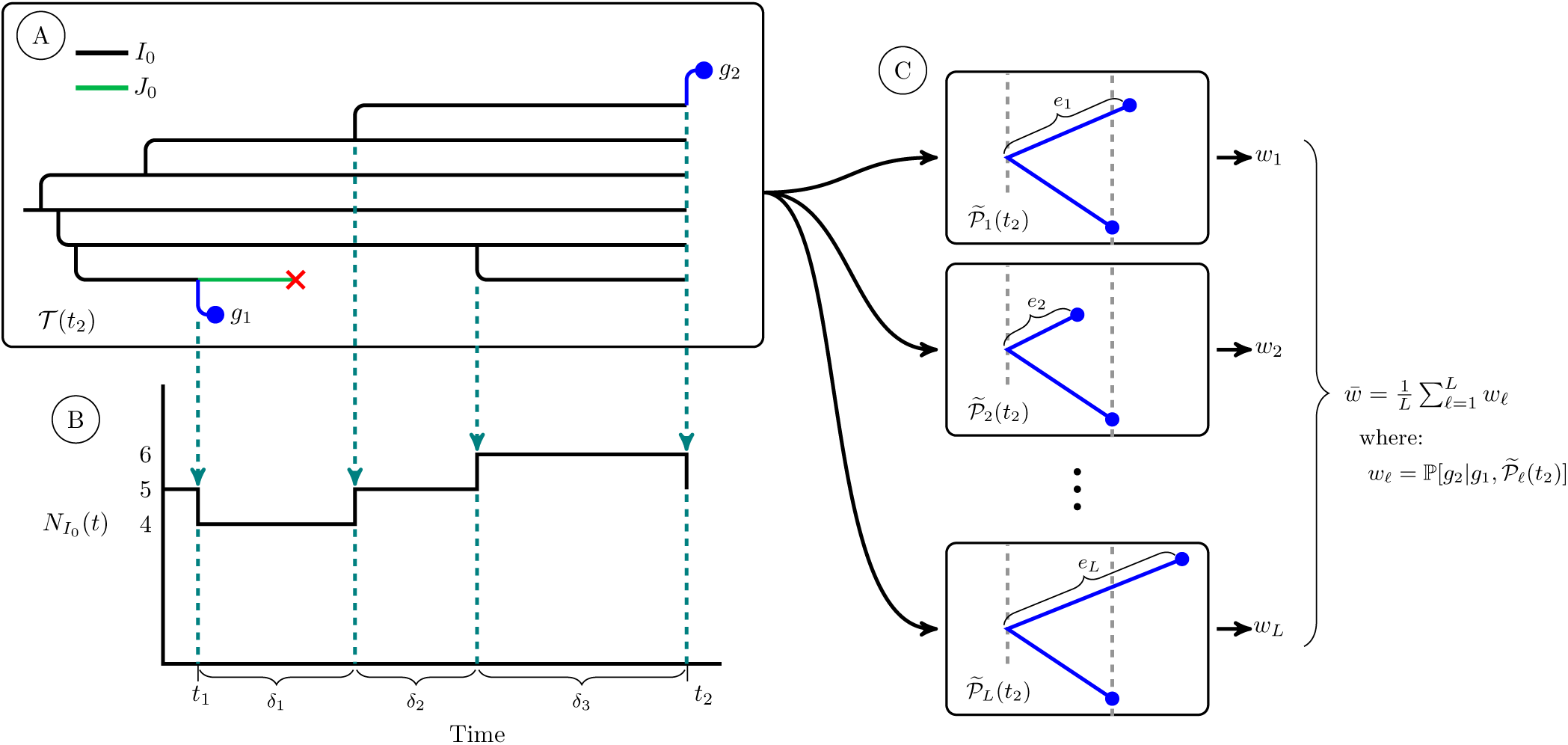
A schematic of quantities used in calculation of the conditional density of a diagnosis and the conditional probability of a genetic sequence. At Ⓐ we show a simulated transmission tree. For simplicity, this tree only has individuals of class *I*_0_ and class *J*_0_. Dashed arrows fall from events in the transmission tree that change the count of *I*_0_ individuals in the population. At Ⓑ we show a plot of the trajectory of the *I*_0_ class. This plot shows the quantities we use to calculate the integrated hazard of diagnosis for the *I*_0_ class, Λ_0_, over an interval of time from *t*_1_ to *t*_2_. We first subdivide the time interval into *R* subintervals over which the number of *I*_0_ individuals is constant (indicated with dashed lines). We let the number of *I*_0_ individuals in the *r*^th^ subinterval be *N*_*I*0,r_. The integrated hazard of diagnosis is then: Λ_0_ = *ρ*_0_ 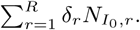 The integrated hazards of diagnosis for the other two classes of undiagnosed individuals are computed in the same fashion. At Ⓒ we show the set of *L* subtrees of the phylogeny that we use to numerically estimate the conditional probability of sequence *g*_2_ under our relaxed clock model. The *l^th^* subtree is constructed by augmenting 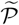(*t*_1_) with a new edge with length *e_l_* drawn from a gamma distribution parameterized as described in the text. For each of these *L* subtrees we use the peeling algorithm to compute 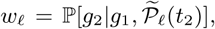 the conditional probability of observing sequence *g*_2_ given sequence *g*_1_ and the structure of 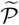_*l*_(*t*_2_). The average of these conditional probabilities is a numerical estimate of the conditional probability of *g*_2_ under our relaxed clock model. For simplicity, here we do not show the case in which the edge length of *g*_2_ splits an existing edge; this case requires a beta bridge to apportion the length of the split edge so as to maintain gamma distributed edge lengths. For this more complicated case, see Fig. 9.

The conditional probability of a genetic sequence is the probability of observing that sequence given the latent state of the system and all previously observed sequences. Our Monte Carlo approach for computing this probability under a relaxed clock is detailed in the ‘Computational Structure’ section.

### 6.5 Data analysis methods: the sequence data

We preprocessed the sequence data following Volz *et al*. (2013a) to facilitate comparision with that work. We excluded poor quality sequences and recombinant sequences, and accounted for known sources of selection. We first aligned all sequences to the reference sequence for the pol gene of HIV subtype-B. We then masked known drug resistant sites, as specified in the Stanford database of HIV drug resistance Bennett *et al*. (2009). We used the program HyPhy Pond *et al*. (2005) to identify the type of each sequence and then excluded recombinant sequences and nonsubtype-B sequences. Many individuals in the dataset have multiple sequences. To limit the complexity of the problem, we chose to keep only first available sequences that were collected within one year of diagnosis. Our methods could, in principle, allow for multiple sequences from each individual. However, this extension has not yet been implemented. We took the time of diagnosis as the time of sequencing – for most sequences this is a reasonable approximation. Poor quality sequencing often manifests as sequences with clipped ends. We therefore considered the length of a sequence as a proxy for quality, and we excluded sequences whose concatenated length was shorter than 1100 base pairs.

### 6.6 Data analysis methods: selecting a subepidemic

The Michigan Department of Community Health (MDCH) maintains an extensive dataset on HIV positive individuals living in the state of Michigan. This dataset stretches back to the beginnings of the HIV epidemic in the United States, and includes over 30,000 diagnoses and nearly 9,000 genetic sequences. Analysis of the full dataset is beyond the scope of our current implementation. Further developments, possibly including preliminary splitting of the full phylogeny into clusters, will be necessary to apply our methods to larger-scale situations. We therefore selected a subset of the cases based on a number of clinical covariates. We chose to focus on the young, black, MSM, subepidemic, which has been of recent concern in Detroit and elsewhere in the USA (Maulsby *et al*., 2014). In selecting this subset, one of our goals was to choose a well-defined subpopulation. We selected records of individuals from the the MDCH dataset that met the following criteria: black, MSM, known not to be an intravenous drug user, and diagnosed in one of 10 counties that comprise the Detroit Metropolitan Area. For this subpopulation, the distribution of the age at diagnosis through time shows striking patterns. In particular, it there is evidence for cohorts of infected individuals that may be clusters of transmission within the young, black, MSM community. We selected a cohort from this population that may represent such a cluster of transmission: individuals that were between the ages of 19 and 28 inclusive in the year 2011 (a span of 10 years) and were diagnosed between 1 January 1999 and 31 December 2011 (Fig. 5). We selected this particular cohort of individuals because it contains what appears to be a pulse of transmission, and because it coincides with when we have high rates of sampling for the genetic sequence data. Counts of individuals diagnosed between 1 January 1999 and 31 December 2003 were used to determine initial conditions (detailed in the supplement). We fit models to data from 1 January 2004 to 31 December 2011. This portion of the cohort has 709 diagnoses and 253 primary genetic sequences. We subsampled the genetic sequences, randomly selecting 100 sequences to keep in the analysis. For the current implementation of our methodology, and in the context of this HIV model, 100 sequences was around the limit of computational tractability.

## Acknowledgments

Data on the the HIV epidemic in Detroit were provided by the Michigan Department of Community Health under a data sharing agreement that received IRB approval. We acknowledge James Koopman and Mary-Grace Brandt for their help in giving us access to these data and for discussions on HIV epidemiology. We are grateful for the support of the Genome Sciences Training Program at the University of Michigan and the following grants: NSF-DMS 1308919, NIH 1-U54-GM111274-01, and NIH 1-U01-GM-110712-01.

## 7 Supplementary Material

Supplementary materials are available online at Molecular Biology and Evolution (http://www.mbe.oxfordjournals.org/). The supplement provides a formal specification of the class of models described in the New Approaches section, technical details on the algorithms we developed to maximize and evaluate the likelihood of these models, and additional details concerning the data analysis presented in the main paper. The source code for our software implementation of the SMC algorithms is available at https://github.com/kingaa/genpomp (to be archived at https://datadryad.org).

## S1 The GenPOMP model: linking infectious disease dynamics with genetic data

We define a class of models that describes an environment within which our general software implementation can be described. We aim at sufficient generality to represent the breadth of applicability of our methodology and the key methodological innovations, yet including enough details to describe the specific data analysis in the main text.

Data consist of *n*\* genetic sequences of a pathogen. We use a convention that *j*:*k* denotes the arithmetic sequence (*j*, *j* + 1,…, *k*), so that the entire collection of genetic sequence data can be written as

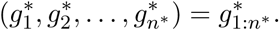

We use asterisks to denote data, to distinguish these from quantities arising in the model. The times at which the sequenced samples are collected are also data, and the total number of sequences, *n*\*, will be modeled as the outcome of a random process rather than some fixed quantity. We write the genetic sequence times as

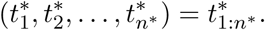

We suppose that the data are collected in a time interval

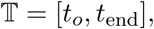

with 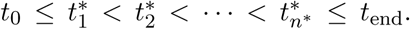 For simplicity, we avoid the possibility of simultaneous observations. If no sequence is available for the diagnosis at some time 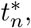 we set 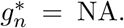 Otherwise, we suppose the collection of sequences 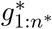 consist of aligned sequences of length *L*, i.e., 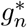 ∈ {*A*,*C*,*T*,*G*}^*L*^.

Here, we do not include the possibility of additional clinical or epidemiological measurements available at diagnosis, though an extension to allow this is fairly straightforward. Further, we consider that only a consensus pathogen sequence is available from each host, so we ignore the possibility of extracting information from data on pathogen genetic diversity within hosts. Nevertheless, our framework can account for sequencing error and differences between observed and transmitted pathogen populations.

The partially observed Markov process (POMP) model consists of a latent, unobservable, Markov process {*X*(*t*),*t* ∈ 𝕋} and an observable process {*Y*(*t*),*t* ∈ 𝕋}. *X*(*t*) takes values in a set 𝕏 and *Y*(*t*) takes values in a set 𝕐. A POMP model for genetic data, which we call a GenPOMP, is required to have the following structure. {*Y*(*t*)} consists of a collection of random number *N* of diagnosis times, denoted *T*_1:*N*_, and corresponding sequences *G*_1:*N*_. The observed outcomes are *N* = *n*\* and (*T_n_*,*G_n_*) = 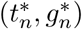 for *n* ∈ 1:*n*\*. We adhere to a convention that random variables are denoted by upper case letters; the corresponding lower case letters are used for possible values of the random variable, and asterisks denote the actual data for observable variables; blackboard bold typeface is used for sets.

Recall that, in the main text, we wrote *X* (*t*) = (𝒯(*t*), 𝒫(*t*), 𝒰(*t*)) where 𝒫(*t*) is a *transmission forest* and 𝒫(*t*) is a *phylogeny*. Here, it is convenient to take a different, but functionally equivalent, approach. We do not require that *X*(*t*) itself contains 𝒯(*t*) and 𝒫(*t*), but we do require that {*X*(*u*), *t*_0_ ≤ *u* ≤ *t*} is sufficient to construct 𝒯(*t*) and 𝒫(*t*). This additional layer of abstraction lets us define the GenPOMP model without having to explicitly construct the processes {𝒯(*t*),*t* ∈ 𝕋} and {𝒫(*t*), *t* ∈ 𝕋}.

The set 𝕏 should describe the state of each individual in a study population. The study population is supposed to contain a finite number of individuals drawn from a countable collection of individuals who could potentially enter the study population. We suppose these potential individuals are labeled with values in the natural numbers, ℕ = {1, 2,3,… }, and so collections of individuals in the study population take values in the set ℍ consisting of all finite subsets of ℕ. We suppose there is a random process {*H*(*t*),*t* ∈ 𝕋}, with *H*(*t*) taking a value in ℍ corresponding to the identities of all individuals in the study population at time *t*. Formally, we suppose that *H*(*t*) is constructed from *X*(*t*) via a suitable function mapping 𝕏 to ℍ. We suppose that each individual in the study population has a state in a set 𝕊. For a simple compartment model, 𝕊 could be finite or countable, however, we also allow for the possibility of continuous real-valued state variables. In particular, we will later define a random clock process governing the rate of pathogen evolution within each individual infected host. To keep track of the state of each member of the study population, we suppose that the state of any individual *i* in the study population at time *t* is given by a random variable *X*_i_(*t*), constructed from *X*(*t*) via a suitable function mapping 𝕏 to 𝕊. A canonical way to do this is to take

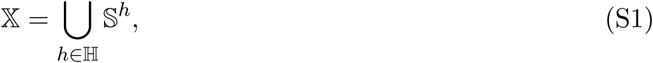

for which an element (*s*_*i*1_, *s*_*i*2_, …,*s*_*i*__*k*_) ∈ 𝕏 is interpreted to mean that the study population is {*i*_1_, *i*_2_,…, *i_k_*} ∈ ℍ and individual *i_j_* is in state *S_j_* ∈ 𝕊. Our definition of the study population is the collection of individuals being modeled, and so the state of individuals outside the study population is necessarily undefined. In order to define {*X_i_*(*t*),*t* ∈ 𝕋} as a stochastic process, one can formally define an additional state ⊙ and set *X_i_*(*t*) = ⊙ when *i* ∉ *H*(*t*). Note that, in general, {*X_i_*(*t*),*t* ∈ 𝕋} does not inherit the Markov property from {*X*(*t*), *t* ∈ 𝕋}. If individual state transitions occur as an independent Markovian process once that individual is infected (as is the case in our HIV example) then {*X_i_*(*t*),*t* ∈ [*t_i_*, *t*_end_]} has a conditional Markov property given *i* ∈ *H*(*t_i_*).

The state process may, in general, need to include other components in addition to {*X_i_*(*t*),*i* ∈ *H*(*t*)}. For example, *X*(*t*) may include dynamic variables affecting the entire population, such as environmental or sociological processes. For the remainder of this article, the specific construction in equation (S1) suffices, but that is not essential to our approach. If 𝕊 is countable then 𝕏, given by (S1), is also countable and {*X*(*t*)} is a Markov chain. Otherwise, {*X*(*t*)} is a more general Markov process.

Some basic properties of individuals characterize the model as a disease transmission system, and these are required to construct the evolutionary process model for the pathogen. This leads us to define functions that return properties about the state of an individual, and we call these *query functions*. This notation differs from usual compartment models, where each individual is modeled as residing in a single compartment. We write properties as functions of *X*(*t*), rather than components of *X*(*t*), to keep applicability to a broad class of population models. As long as the required query functions can be defined for a population model, the statistical methodology developed will apply, giving the scientist considerable flexibility in the specification of the model.

We require that an individual’s state, i.e., its value in 𝕊, can describe whether that individual is infected and infectious. We represent this requirement by supposing that there is a query function

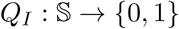

defined as,

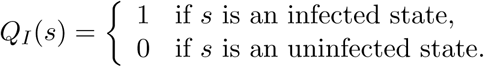

To link the model to diagnosis data, we require that a state in 𝕊 determines whether an individual is diagnosed while part of the study population. Specifically, we suppose there is a query function

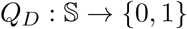

such that

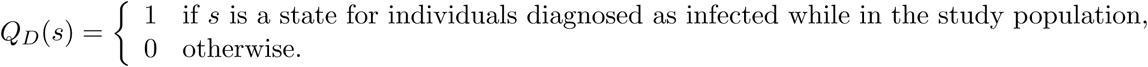

We then define

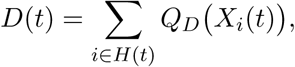

which counts the number of individuals diagnosed while in the study population, by time *t*. This counting process (i.e., a non-decreasing integer-valued process) is relevant for relating the model to the data on the study population. Note that *D*(*t*) does not count the number of clinically diagnosed individuals in the study population at time *t*, which would require a different accounting for the possibility of immigration and emigration of diagnosed individuals.

Now, we define the set of infected states to be

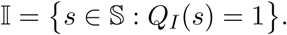

We suppose that the state contains information about the identify of the infector, and we do this by requiring the existence of a query function

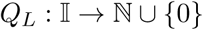

defined such that

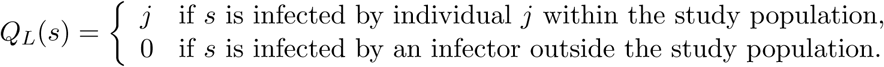

The capability to construct the query function *Q_L_*(*s*) requires that the identity of the infector is stored in the state variable at the point of infection, so it is available later as part of the state of the infectee. Information on the identity of the infector is not usually required for a compartment model, but is useful when working with genetic data in order to track lineages of the pathogen.

The evolutionary process of the pathogen genome within an individual in the host populations is modeled using a relaxed molecular clock, meaning that standard molecular models for evolution are applied on a stochastically perturbed timescale. It has become established that the usual models for molecular evolution fit sequence data better if one allows such fluctuations in the rate of evolution (Drummond et al., 2006). To implement a relaxed clock, we construct a random process on each edge of the transmission tree. This process scales calendar time to evolutionary time, the latter meaning a modified timescale on which the evolutionary rate is constant. We therefore require the existence of a query function

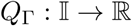

returning the relaxed evolutionary clock time corresponding to evolution of a transmissible pathogen population within an infected individual. Specifically, *Q*_Γ_(*s*) represents the random, individual-specific, clock time for the evolutionary process that separates the host’s transmissible pathogen population from the rest of the pathogen community when the host is in state *s* ∈ 𝕀. For an individual based model in which an individual is infected within the study population, this corresponds to the evolutionary process within the host subsequent to infection. Immigrant pathogens require additional assumptions on how they relate genetically to pathogens already circulating in the study population. Conditional on the randomly perturbed molecular clock, pathogen evolution is usually specified by a general time-reversible Markov model.

We also suppose the existence of a query function

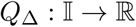

which returns the relaxed evolutionary clock time separating the measurable pathogen population from the transmissible host population within an infected individual. If and when an individual gives rise to a pathogen genetic sequence within the dataset, this clock time adds to the clock time *Q*_Γ_(*s*) in determining the probability distribution of the measured sequence.

The separation of the pathogen evolutionary process into transmitted and untransmitted mutations has multiple interpretations. The choice of primary interpretation has consequences for the appropriate model specification of the branch separating the measurement node v from the transmission tree. The plausibility of these different interpretations will depend on the biological system under investigation.

(B1) Measurement error. Sequencing error could be modeled by an arbitrary evolution-like process on the branch separating the measured sequence from the transmissible sequence.

(B2) Transmissible versus measurable strains. The measured sequence may reflect the dominant strain reproducing most competitively within the host. It is conceivable that much of the diversity resulting from within-host evolution may lead to pathogens which are non-viable or non-competitive for between-host transmission. The evolutionary branch corresponding to the measurement event could represent this dead-end evolution, leaving the main body of the transmission tree to represent evolution of a transmissible strain.

(B3) Within-host diversity. A strain transmitted subsequent to sequencing could be more similar to an ancestral strain than to the sequenced strain by chance, due to within-host genetic variation, even without appealing to a phenomenon such as (B2). The measurement branch permits such behavior, so may help to adjust for unmodeled within-host pathogen genetic diversity.

Other model-specific quantities can be defined by additional query functions, but are not essential components of a GenPOMP model. For example, epidemiological models commonly consider the number of susceptible or removed individuals. Also, having defined an appropriate query function for a category of individuals, one can define a process counting such individuals. For example, to complement the query function *Q_I_* for infected individuals, we can define a process

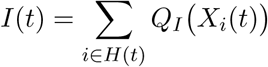

counting the number of infected individuals in the study population. We can also write the size of the study population at time *t* as,

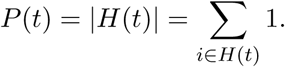

Our framework therefore incorporates the structure of arbitrary compartment models (Breto et al., 2009) represented at the level of compartment membership for each individual.

The history of the query functions for infected individuals,

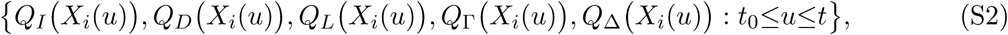

is sufficient to construct the transmission forest, 𝒯(*t*), and phylogeny, 𝒫(*t*), described in the New Approaches section of the main text. Formally, for (S2) and elsewhere, we extend the query functions to take an undefined value, denoted by ⊙, when the argument is outside the defined domain. To specify the measurement process model, recall that the measurement process {*Y*(*t*)} consists of an increasing sequence of diagnosis times {*T_n_*} associated with the diagnosis counting process {*D*(*t*)}, together with a collection of genetic sequences {*G_n_*}. We suppose that the sequences {*G_n_*} are modeled as a continuous-time Markov chain on 𝒫(*t*). The probability distribution of the genetic sequence *G_n_* at time *T_n_*, conditional on {*X*(*t*),*t* ≤ *t_n_*} and *G*_1:*n*–1_, therefore depends on 𝒫(*t*) and *G*_1:*n*–1_. If a genetic sequence for the diagnosis at time *T_n_* is not available, we assign *G_n_* the value NA. We suppose this occurs with probability 1 – *pG*, independently of {*X*(*t*)}.

We have defined the GenPOMP model so that the pathogen genetic sequence arises only in the measurement model. No genetic sequences are included in the state process, or its particle representation. Our approach is consistent with viewing the genetic evolutionary model as a principled way to define and evaluate a statistical metric between genetic sequences that respects the tree structure of the evolutionary process and has the property that similar sequences are more likely to come from closely related pathogens. A measurement model satisfying these criteria and providing a statistical fit to the data need not be judged on the details of its biological strengths and weaknesses if the microevolutionary processes are not the focus of the investigation. The individual, stochastic molecular clocks determining the rate of evolution within each host are included in the latent process component of the GenPOMP model to facilitate Monte Carlo integration over the distribution of these clocks, as described in Section S2.

The definition of a GenPOMP model given here is general and abstract. The population model {*X*(*t*) } corresponds to an arbitrary individual-based Markovian model constrained to include concepts of transmission of a pathogen and measurement of pathogen genetic sequences. The measurement model is constrained to be based on a Markovian evolutionary process, but this is standard in current models used for phylodynamic inference. Our methodological approach applies to this general GenPOMP model class, subject to being able to simulate from the individual-based model and compute the rate at which individual hosts provide a pathogen sequence. The Markovian assumption is convenient algorithmically. In one sense, it is not fundamentally a limitation since non-Markovian models may be approximated by Markovian models with additional state variables. In another sense, it is a practical limitation since increasing the size of the state space increased the computational effort required.

## S2 A GenSMC algorithm for filtering the GenPOMP model

We develop a sequential Monte Carlo (SMC) approach for the framework of Section S1. We will use the name GenSMC to describe an SMC algorithm for GenPOMP models. As an instance of SMC, the basic principles and theoretical foundation for GenSMC follows from the general theory of SMC (Liu, 2001). However, GenPOMP models have a particular structure that places particular demands on a GenSMC algorithm. Many variations are possible on our GenSMC algorithm, but demonstration of one successful GenSMC algorithm provides a foundation and motivation for future improvements. Our GenSMC approach is presented as pseudocode in Algorithm S-1, which is an expanded version of Algorithm 1 in the main text. We proceed to define the notation that will be required.

To construct our algorithm, we specify a concrete class of GenPOMP models. Let {*X*(*t*),*t* ∈ 𝕋} be a latent GenPOMP process with the form

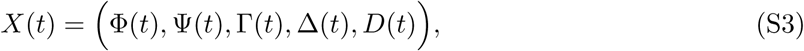

having components Φ(*t*), Φ(*t*), Γ(*t*), Δ(*t*)and *D*(*t*) defined as follows:

{*D*(*t*)} records diagnosis events within the study population, as defined in Section S1. We suppose that no diagnoses occur simultaneously, so {*D*(*t*)} is a *simple* counting process. Therefore, we can model {*D*(*t*)} via a conditional intensity process *ρ*(Φ(*t*)), Ψ(*t*) such that

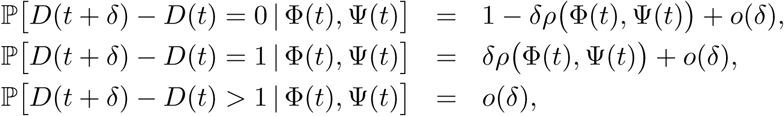

where *o*(*δ*) denotes a function *f* : [0, ∞) → ℝ satisfying lim_*δ*→0_ *f* (*δ*)/*δ* = 0. Here, *ρ*(*X*(*t*)) is called the diagnosis rate.

{Ψ(*t*)} is a piecewise constant process which records a list of the identity labels of individuals diagnosed by time *t*.

{Φ(*t*)} contains everything else in the GenPOMP model, so is essentially arbitrary within the general requirements of a GenPOMP model. We suppose that observation events are also recorded in the state process; specifically, the observation counting process {*D*(*t*)} is a function of {Φ(*t*)} which gives rise to observation times {*T*_1_,*T*_2_,… } at which the genetic measurements {*G*_1_, *G*_2_,… } are made.

{Γ(*t*)} is a list of the relaxed clock process for all the interior edges of the transmission tree, i.e., Γ(*t*) = {*Q*_Γ_(*X_i_*(*t*)),*i* ∈ ℕ} where *Q*_Γ_ is defined in Section S1.

{Δ(*t*)} is a list of the relaxed clock process for the terminal branches of the transmission tree,i. e., Δ(*t*) = {*Q*_Δ_(*X_i_*(*t*)), *i* ∈ ℕ} where *Q*_Δ_ is defined in Section S1.

The relaxed clock processes affect the micro-evolution of the pathogen, but in our model the genetic process has no consequence for the transmission dynamics: the genetic sequence is simply a marker, and the genetic models we use are models for neutral evolution. A consequence of this is that the relaxed clock processes only have to be evaluated when needed to compute the conditional probability mass function for attaching a new genetic sequence to the genetic tree. If these components of the latent process can be computed when needed, there is no need to continually update them. Our computational strategy to take advantage of this is called a just-in-time representation and is formally described in Section S3.4. Informally, the just-in-time representation is the tool that lets us define the latent GenPOMP model as a continuous-time Markov process while updating the relaxed clock processes at diagnosis times, when needed. To simulate the GenPOMP model forward in time using a just-in-time representation, we need to be able to evaluate the relaxed clock process over arbitrary time intervals, and also split the evolutionary time over a branch of the transmission tree if a new measurement divides this branch. An example of a Markovian clock with these properties is the Gamma process.

We will show that the relaxed clock processes {Γ(*t*)} and {Δ(*t*)} can be represented by two processes {*U*(*t*)} and {*V*(*t*)} which generate the evolutionary clocks that are necessary to evaluate the likelihood of the sequences. The processes {*U*(*t*)} and {*V*(*t*)} are constant except at diagnosis times, and so are fully specified by the discrete processes *U*_0:*N*_ and *V*_0:*N*_, with *U_n_* = *U*(*T_n_*) and *V_n_* = *V*(*T_n_*). The construction of {*U*(*t*), *V*(*t*)} is an instance of just-in-time variables, as discussed further in Section S3.4. Therefore, for the purposes of Algorithm S-1, it is convenient to replace the representation in equation (S3) with an equivalent representation,

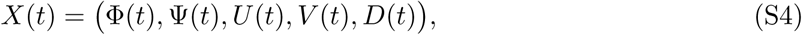

The construction of {*U*(*t*)} and {*V*(*t*)} is described in Figure S-1

Algorithm S-1 is written using discrete time steps corresponding to the sequence of observation times, together with the start and end times of the interval 𝕋. It convenient to define

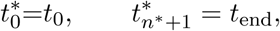

so that 𝕋 = 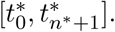 {Ψ(*t*)} is fully specified by its values at the discrete set of observation times, and so we define a process {Ψ_*n*_} with

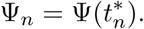

**Figure S-1:**
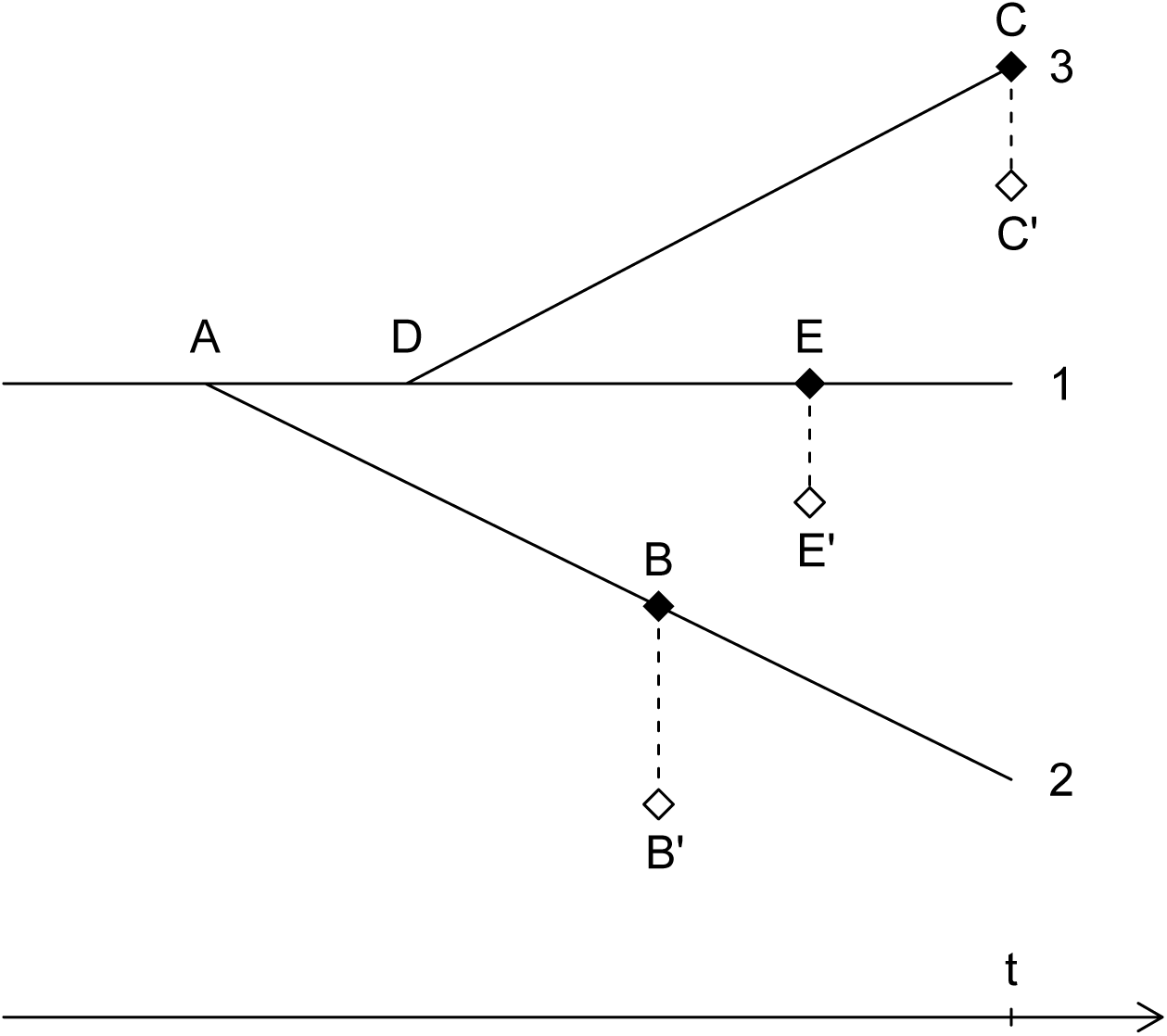
The diagram represents the transmission tree for a particle where individual 1 infected individual 2 at time *A* < *t* and individual 3 at time *D* < *t*. Sequences are collected at times *B*, *C* and *E*. Measured but untransmitted sequence mutations occur along *BB′*, *CC′* and *EE′*. For this particle, we know that the sequence at time *B* corresponds to individual 2, and the sequence at time *E* belongs to individual 1. Suppose we then wish to evaluate the probability of the new sequence at time *t* conditional on it belonging to individual 3, as shown on the diagram. From the previous observed sequences, assigned to *B′* and *E′*, this particle has already been assigned evolutionary clock times for the segments *AB′* and *AE′*. To place the new sequence at *C′*, we first generate a new clock process for the segment *DC′*, which is represented by the variable 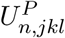 in step 8 of Algorithm S-1. Then, we split the evolutionary clock time for *AE′* into *AD* and *DE′*, in a way that is consistent with the corresponding calendar times and the stochastic evolutionary clock process. This computation is represented by the variable 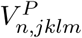 in step 10 of Algorithm S-1.

To provide a discrete time representation of {Φ(*t*)}, we write

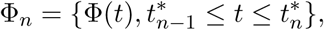

for *n* = 1,…, *n*\* + 1, with Φ_0_ = Φ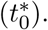 Similarly, we write

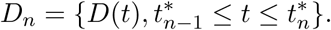

Diagnosis events are modeled as perfectly observed, almost tautologically. We write *d*\*(*t*) for the observed value of *D*(*t*), defined as

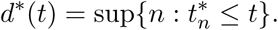

Also, we write 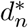 for the observed value of *D_n_*. Perfectly observed components of the latent process of a POMP model require special attention in sequential Monte Carlo algorithms, and so Algorithm S-1 uses the targeted proposal developed in Section S3.2 to handle the diagnosis process.

Hierarchical sampling (described in Section S3.3) is carried out in Algorithm S-1 over the components Φ(*t*) and Ψ(*t*) in (S3), as well as over the components *U_n_* and *V_n_* in the just-in-time representation of {Γ(*t*)} and {Δ(*t*)}.

The pseudocode for Algorithm S-1 adopts a space-saving convention that index *j* always ranges over 1 : *J*, index *k* ranges over 1 : *K*, index *l* ranges over 1 : *L*, and index *m* ranges over 1 : *M*. Thus, for example, line 6 of Algorithm S-1 has an implicit loop over *j* ∈ 1 : *J* and *k* ∈ 1 : *K*.

If 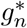 = NA then *w*_2_(*n*, *j*,*k*,*l*, *m*) is defined to be the probability of not recording a genetic sequence at diagnosis. In this case, steps 7 to 16 are not necessary: it suffices to take *K* = 1, with *U_n_* and *V_n_* being undefined. This special case is omitted from Algorithm S-1 for simplicity.

To implement Algorithm S-1, we require code to generate initial values, and to simulate the dynamic model for all the hierarchical layers conditional on the diagnosis events. Specifically, we require simulators for

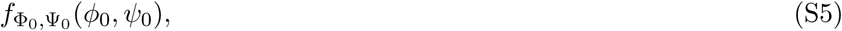

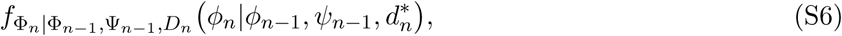

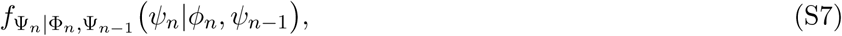

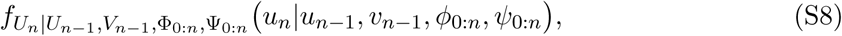

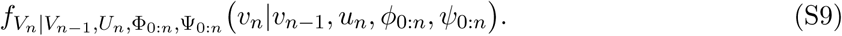

We then require code to evaluate the diagnosis rate,

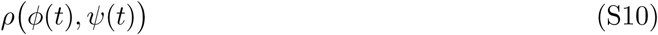

as well as the genetic measurement model,

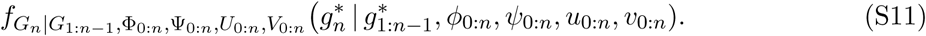

All the densities in (S5-S11) may additionally depend on a parameter vector *θ*.

##### Algorithm S-1: GenSMC

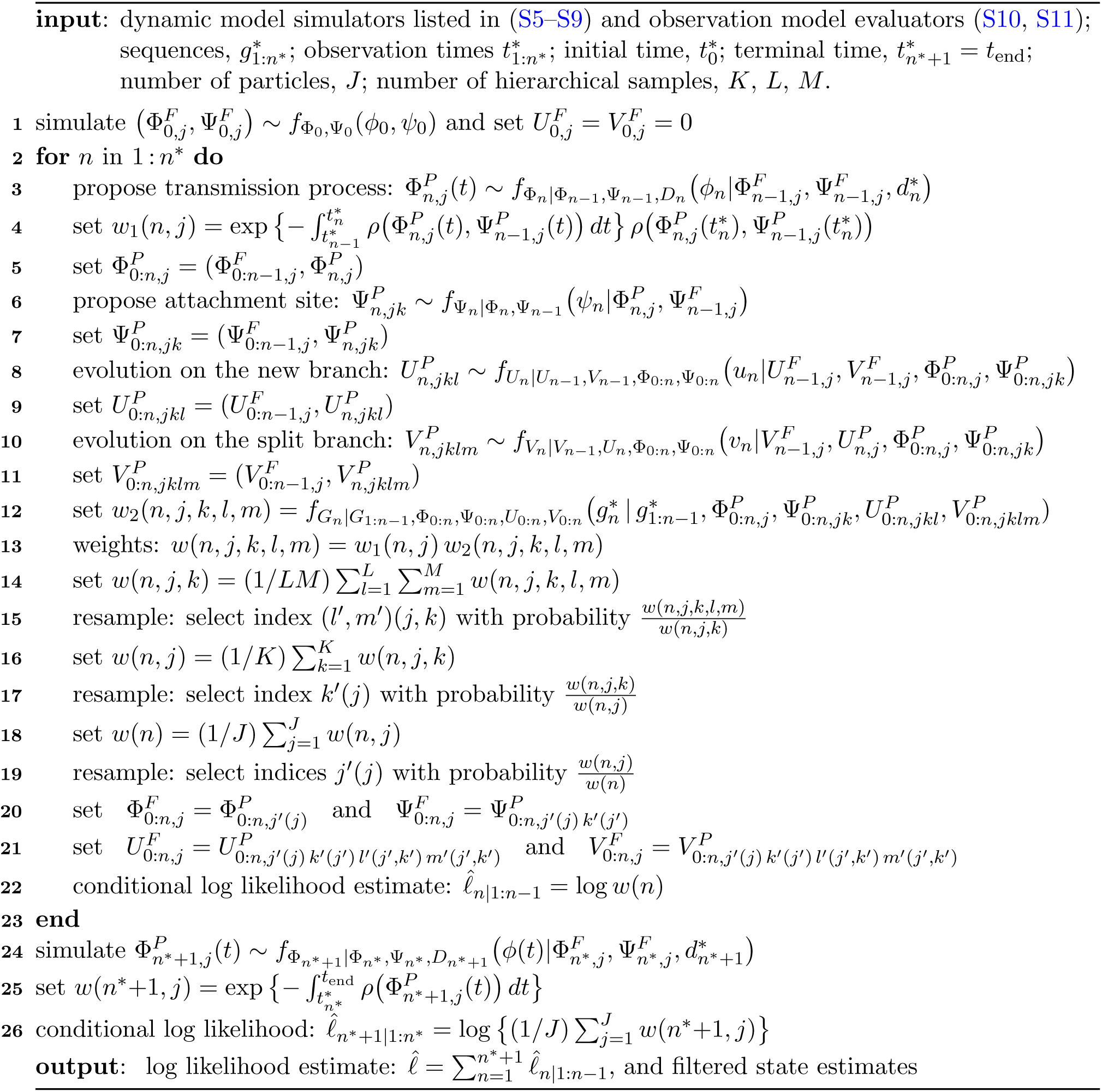

### S2.1 The implementation of GenSMC in the genPomp program

Many computational issues arise in effective implementation of a GenSMC method such as Algorithm S-1. Data structures are needed to keep track of the individuals in the study population, and the genetic relationships between pathogens in different hosts. Efficient implementation of all these computations, including use of a multi-processor computing environment, is necessary to work on problems of a practical scientific scale. The record of our implementation is the open-souce code for the genPomp program that we developed to carry out inference for GenPOMP models, available at https://github.com/kingaa/genpomp. The accuracy of `genPomp` has been successfully tested against exact analytic calculations for some very small scale situations, and against the pomp package (King et al., 2016) for situations where no diagnoses lead to genetic sequences.

There is a substantial difference in the level of abstraction between the formal mathematical representation of a GenPOMP model in Algorithm S-1 and the practical implementation in `genPomp`. One could write more pseudocode to bridge this gap, but that is beyond the scope of this article. We have focused instead on the foundational task of understanding how Algorithm S-1 fits in with the theory and practice of SMC.

### S2.2 Extending GenSMC to infer unknown parameters: The GenIF algorithm

Sequential Monte Carlo (SMC) algorithms such as Algorithm S-1 produce a Monte Carlo approximation to the likelihood of the model, but do not directly provide estimates of unknown parameters. A substantial literature has emerged on using SMC as a basis for statistical inference (Kantas et al., 2015). Iterated filtering (Ionides et al., 2006, 2015) uses SMC, together with parameter perturbations, to maximize the likelihood function. Iterated filtering has demonstrated effectiveness on various nonlinear models arising in infectious disease transmission studies (Ionides et al., 2015, and references therein). We developed an adaptation of Algorithm IF2 of Ionides et al. (2015), which we call GenIF as an abbreviation of *iterated filtering for GenPOMP models*. Our implementation of this GenIF algorithm is included within the `genPomp` program, as described fully in the source code. Conceptually, and computationally, GenIF is a simple extension to GenSMC. GenIF carries out multiple iterations of Algorithm S-1 (GenSMC) adding perturbations to the candidate values of unknown parameters. GenSMC selects particles consistent with the data, and so allowing particles to have diversity in their parameters values naturally selects for parameter values consistent with the data. The theory and practice of iterated filtering focuses on using this phenomenon, with multiple SMC iterations having perturbations of decreasing magnitude, to maximize the likelihood. Previous iterated filtering theory does not encompass the just-in-time variables employed by GenSMC. In the context of GenIF, this means that the current theoretical justification of IF2 (Ionides et al., 2015) does not perfectly apply when we carry out inference for the molecular evolution parameters. Heuristically, however, the principle of iterated filtering still applies, and we rely on empirical results to confirm that maximization performance is satisfactory.

Algorithms that permit numerically satisfactory likelihood maximization and likelihood evaluation provide a foundation for carrying out likelihood-based statistical inference. Profile likelihood methods can be used for obtaining confidence intervals, and likelihood ratio tests or Akaike’s information criterion can be used for model selection.

### S2.3 Scalability of GenSMC

**Figure S-2:**
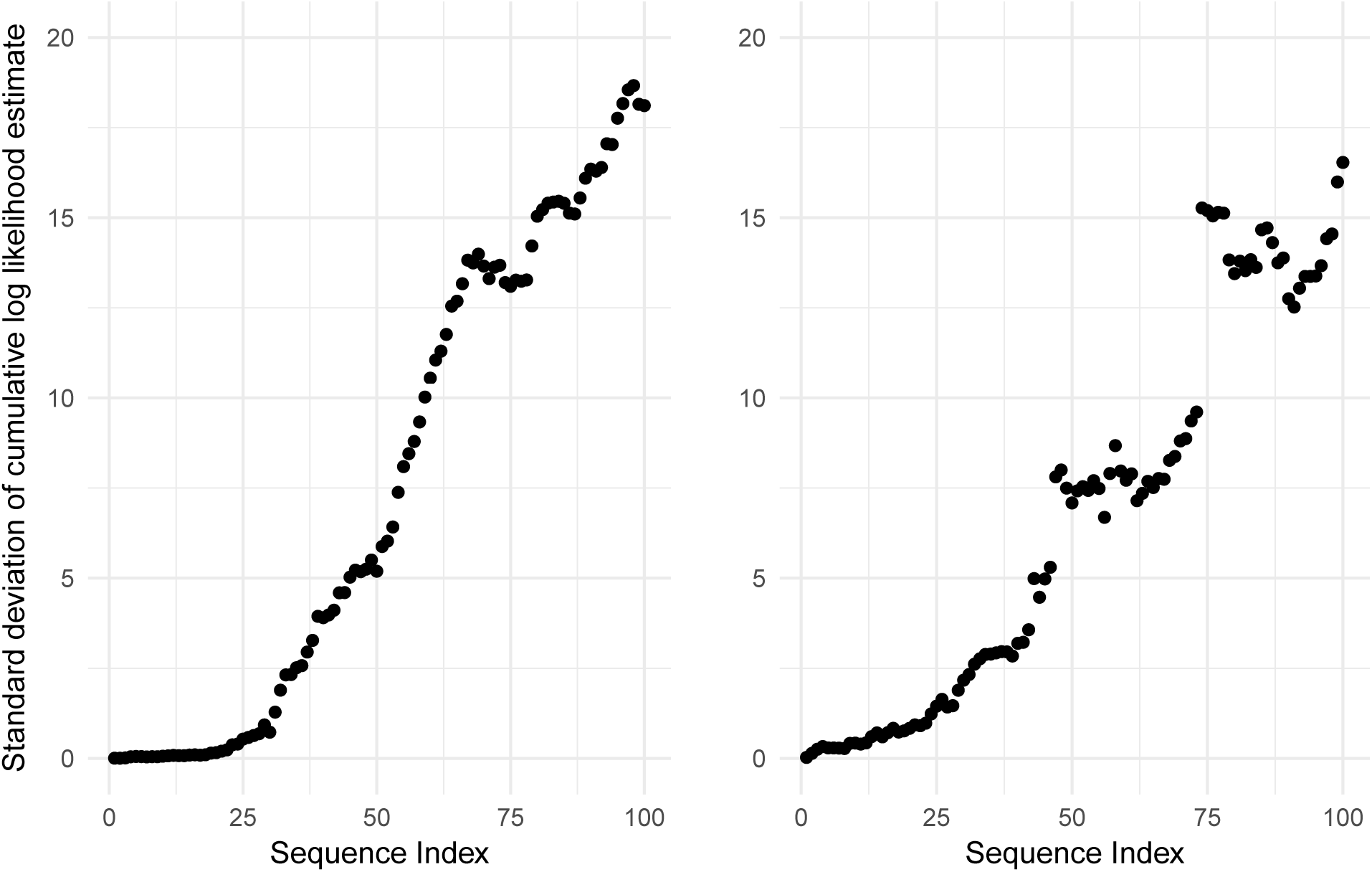
Results from two experiments exploring how the standard deviation in the log likelihood estimate scales with the number of sequences fit. On the left, results from fitting to simulated sequences. On the right, results from fits at the MLE in the data analysis.

To explore the scalability of our GenSMC implementation, we performed two experiments: one with simulated data and one with real data. For the simulated data, we first simulated an epidemic conditional on observing 100 sequences. We then ran the particle filter at the truth 10 times, each time using 10,000 particles. Finally, for each sequence, we computed the standard deviation of the cumulative log likelihood estimate across the 10 filtering evaluations. This computation yields a measure of the variability in the log likelihood estimate if one were to stop filtering at each sequence. For the real data, we performed the same calculation using filtering results from fits at the MLE of the data analysis, again using 10 particle filters each with 10,000 particles. The results from simulated data provide a controlled assessment of how Monte Carlo variance scales as the number of sequences grows. The results from real data give us one example of how Monte Carlo variance scales in practice. In both cases, the standard deviation of the log likelihood estimate remains relatively small up to around 25 sequences (Figure S-2). An interpretation of this is that placing early sequences on the growing phylogenetic tree is relatively easy. It can become harder to find trees with appropriate places to attach later sequences, leading to increasing Monte Carlo variance. Monte Carlo variance is expected to grow as the size of a computational problem increases, but we did not find a rapid exponential growth. The peeling algorithm for computing the likelihood of the genetic sequences conditional on the phylogeny was typically the largest computational component, though not for all regions of parameter space.

## S3 A theoretical derivation of the GenSMC algorithm

To derive and justify GenSMC (Algorithm S-1) for the GenPOMP model, we work up in stages from a simple and standard SMC algorithm. Initially working in discrete time, we start in Section S3.1 by writing an SMC algorithm that allows for general dependence between the latent process and the observation process. Then, we consider a useful class of targeted proposal distributions in Section S3.2. We add hierarchical layers of resampling in Section S3.3. In Section S3.4, we consider a *just-in-time* approach to construction of state variables which can have their creation postponed until necessary. In Section S3.5, we move these developments into the context of continuous time models. Putting these components together, we obtain Algorithm S-1.

### S3.1 A basic SMC algorithm

Consider a model consisting of a latent stochastic process *X*_0:*N*_ = (*X*_0_, *X*_1_,…, *X_N_*) and an observable process *Y*_1:*N*_ = (*Y*_1_, *Y*_2_,…, *Y*_N_). In this setting, *N* corresponds to the number of discrete time points, differing from the notation of Section S1. Data consist of a sequence 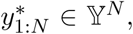 modeled as a realization of *Y*_1:*N*_. We suppose *X_n_* and *Y_n_* take values in measurable spaces 𝕏 and 𝕐, and we require the existence of a joint density *f*_x_0:*N*_,y_1:*N*__ on 𝕏^*N*+1^ × 𝕐^*N*^. Conditional densities are denoted using subscripts, for example, the density of *Y_n_* given *Y*_1:*n*–1_ and *X*_0:*n*_ is written as

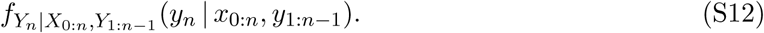

In a standard POMP model, {*X_n_*} is a latent Markov process and the conditional distribution of *Y_n_* depends only on *X_n_* (Bretó et al., 2009). In the context of GenPOMP, we require the marginal Markov property for the latent process,

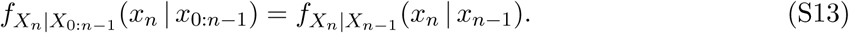

but we allow a general form for the measurement model in equation (S12), where the conditional distribution of the *n*th observation can depend on the entire histories of the latent process and the observation process. SMC techniques for POMP models can be extended to this more general dependence structure (Liu, 2001). A basic SMC algorithm is outlined in Algorithm S-2. This is essentially the basic bootstrap filter algorithm of Gordon et al. (1993), generalized to allow for the dependence on the history of the process in (S12). Notationally, for Algorithm S-2 we set 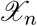 = *X*_0:*n*_ and use superscripts *F* and *P* to denote particles representing the filtering and prediction distributions respectively. We use systematic resampling in place of multinomial resampling (Arulampalam et al., 2002; Douc et al., 2005).

#### Algorithm S-2: A basic Sequential Monte Carlo (SMC) algorithm

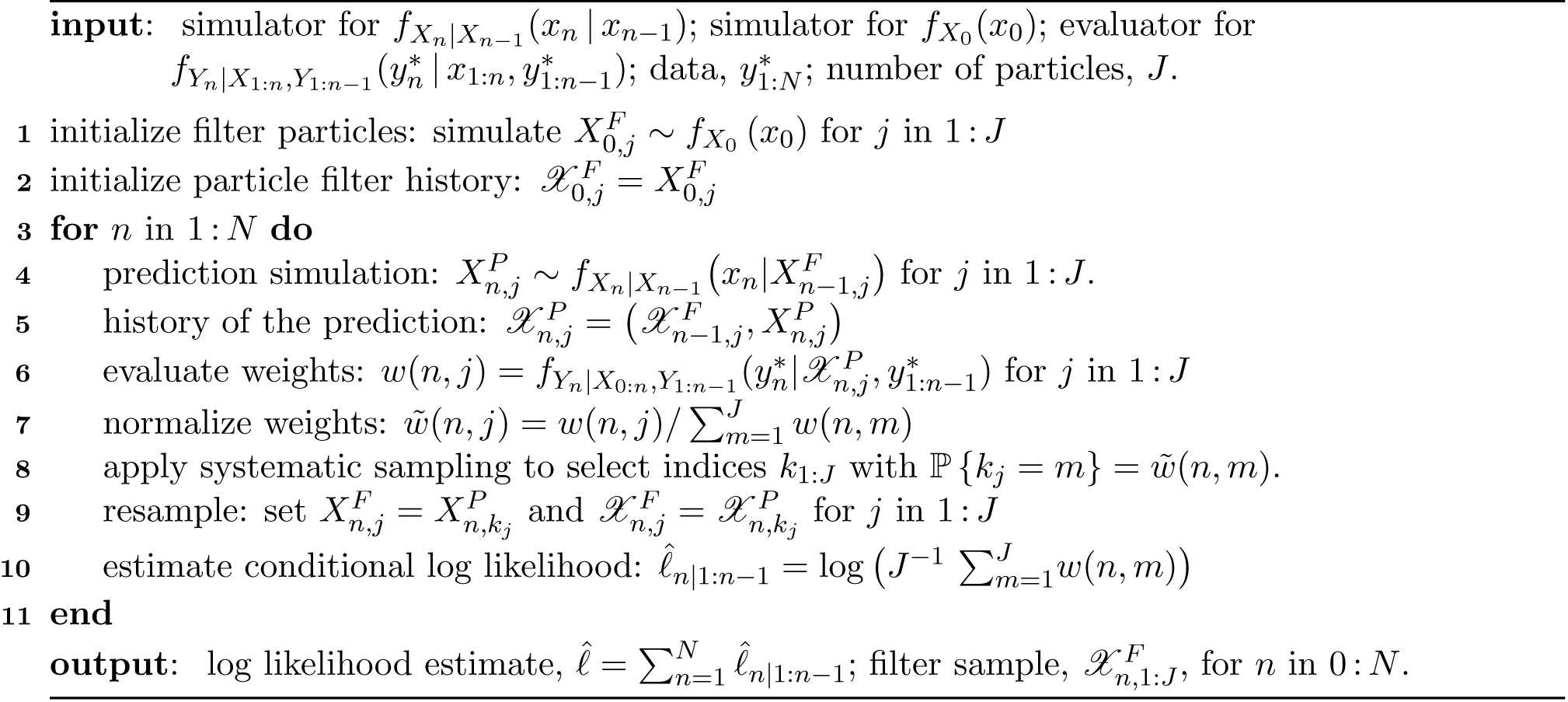

Computational resources are an issue for GenPOMP models, since the spaces 𝕏 and 𝕐 are both large. Furthermore, the dependence on the history in (S12) leads to additional computational requirements for both memory and numerical operations. Careful implementation of SMC is therefore necessary to make the approach practical. We therefore proceed to develop extensions of Algorithm S-2 that are necessary to improve numerical tractability for GenPOMP models.

To understand Algorithm S-2, and subsequently extend it, we write out an algebraic justification of the prediction and filtering steps. For a general latent process *X*_0:*N*_ and observable process *Y*_1:*N*_ modeling data 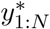 collected at times *t*_1:*N*_, assuming (S12) and (S13), the *prediction identity* is

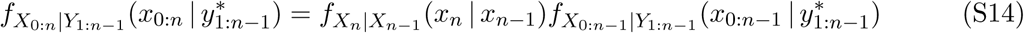

The SMC interpretation of (S14) is that 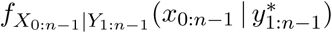 is represented by a collection of of *J* filter particles 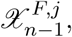 *j* = 1,…, *J*. Algorithm S-2 corresponds to a basic version of SMC in which particle *j* has a time *t_n_* value generated from 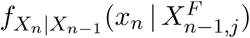 to give rise to a time *t_n_* prediction particle 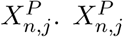 inherits its history from 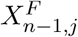 and so 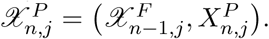

A general *filtering identity* is

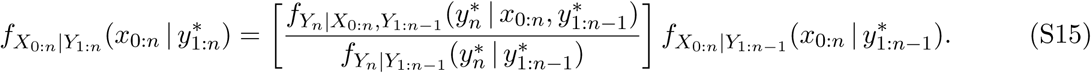

The SMC interpretation of (S15) is that observation 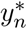 requires the prediction particle 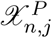 representing 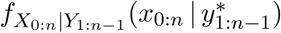 to be given a weight proportional to 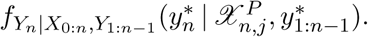 The denominator on the right hand side of (S15) is an irrelevant constant for computing the normalized weights. However, this denominator is approximated in Algorithm S-2 as the normalizing constant, giving a Monte Carlo estimate of the nth term in a factorization of the likelihood of the data,

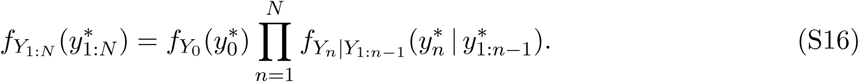

For a discrete time representation of a simple GenPOMP model, Algorithm S-2 might be directly applicable. For example one can take *X_n_* to correspond to all the information about individuals in the population at time *n*, so that *X*_0:*n*_ includes the transmission tree. We could also suppose that *X*_0:*n*_ includes information on who would get sequenced if there are observed sequences—but not how many sequences were observed, which is part of the measurement. For example, at each time point *t_n_*, the state could contain a permutation listing the order in which eligible individuals are sequenced. This construction may appear somewhat contrived, and we proceed to relax it by allowing part of the latent process to be fully observed and therefore also be part of the measurement process. Regardless of that issue, evaluation of 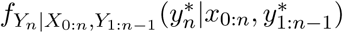 involves evaluating the likelihood of a phylogeny, which can be computed efficiently by a peeling algorithm, together with term for the probability of the sequence being collected.

### S3.2 A targeted SMC approach with a partial plug-and-play property

Some models of interest may have the feature that the event of obtaining a measurement has an appreciable consequence for the latent dynamics. HIV, for example, has the features that sequencing of the pathogen typically occurs at diagnosis. The fraction of infections which are sequenced is high, and diagnosis plays an important role in transmission dynamics both through changes in sexual contact behavior and reduced infectivity due to antiviral drugs. For HIV, it is therefore natural to consider models where sequencing events correspond to transitions of an individual between states and therefore correspond to a perfectly observed component of the latent process. This kind of situation needs some extra care, since 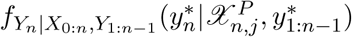 in Algorithm S-2 becomes zero for every draw of 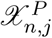 which is not consistent with 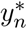. The standard SMC approach to this is to allow for the possibility of targeted SMC proposal distributions, not necessarily the “vanilla” choice *f_Xn_*|*X*_*n*–1_. Suppose the proposal distribution for the SMC algorithm is 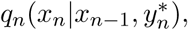 which is permissible since the proposal distribution is in general allowed to depend on any past, current or future observations. This corresponds to rewriting (S14) as

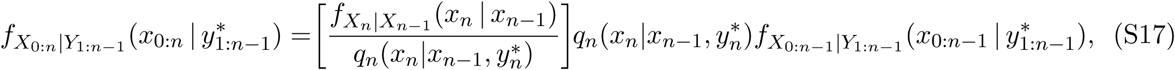

which is interpreted to mean that the targeted SMC proposal particle 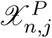, with 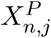 drawn from 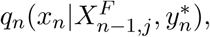 must be given a weight 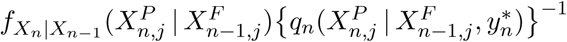 corresponding to the ratio in square brackets in (S17).

A special case of a targeted proposal arises in the situation where part of the state variable is perfectly observed. To describe this situation, suppose we can partition the latent and observable processes as,

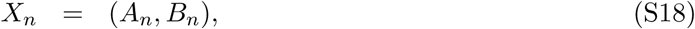

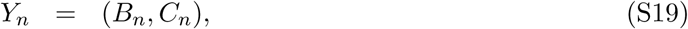

with the data being 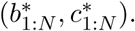 The prediction identity in (S17) can then be written as

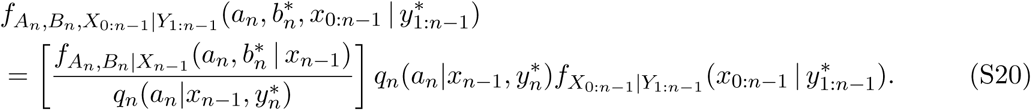

Then, to obtain the filtering distribution 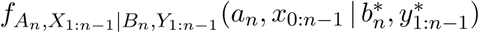 one normalizes the weighted particle representation of 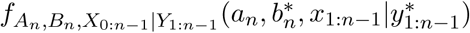 in (S20), with the normalizing constant being the conditional likelihood, 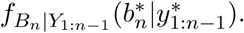 A particular target choice of interest in (S20) is

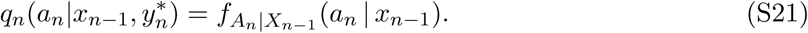

(S20) becomes

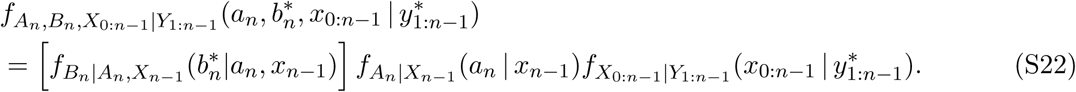

On the component of the state space that is not perfectly observed, the proposal in (S21) is *plug-and-play* (Breto et al., 2009; He et al., 2010) meaning that the algorithm needs only a simulation from *f*_A_n_|*x*_*n*–1__ (*a_n_* | *x*_*n*–1_). However, we require numerically tractable evaluation of the importance sampling weight

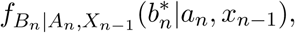

arising from the identity (S22), and so we describe the algorithm as *partially plug and play*.

Using a targeted proposal typically leads to algorithms without the plug-and-play property. Here, we work with situations where 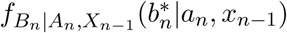 is tractable, even if the complete transition density of (*A_n_*,*B_n_*) is intractable. Thus, *f*_*A*_*n*_ |*X*_*n*–1__ (*a_n_* | *x*_*n*–1_) can be specified in a fairly arbitrary way.

**Example 1**. *B_n_* might be the number of diagnoses at time *n*, which might have a Poisson or negative binomial distribution conditional on *A_n_*.

**Example 2**. Writing the number of sequenced diagnoses at time *n* by 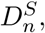 unsequenced diagnoses by 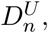 and infected individuals by *I_n_*, we might have *B_n_* = 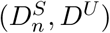 and *A_n_* = *I_n_*. The joint distribution of 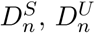 and *I_n_* 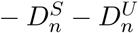 might be multinomial given *I_n_*.

**Example 3**. *B_n_* might describe the race or age group of diagnosed individuals as well as whether they were sequenced.

### S3.3 SMC with hierarchical sampling

For computational considerations, it may be preferable to maintain *J* filtering particles and generate *K* prediction particles from each, rather than maintaining *JK* filtering particles. Computation of the *K* prediction particles can be localized on a single core of multi-processor hardware, and the memory usage may scale with *J* rather than *JK*.

In the context of Algorithm S-2, extended to include the general proposal distribution of Section S3.2, we write 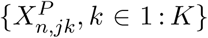 for *K* draws from 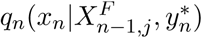 for each value of *j*. We compute the weights in the second layer of the hierarchy by

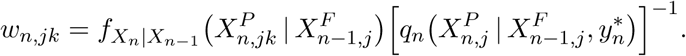

We then define 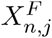 to be a draw from 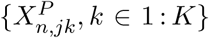 with probability proportional to *w*_*n*,*jk*_, with the history 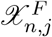 being constructed accordingly. We then assign 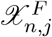 a weight

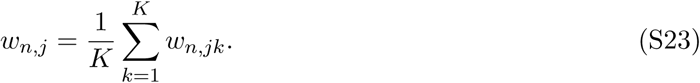

The filter particles {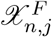, *j* = 1,…, *J*} can be again resampled with weight proportional to *w*_*n*,*j*_ if so desired. Resampling each layer of the hierarchy one at a time gives an approach that we call *staggered resampling*. It might sometimes be preferable to resample *J* particles from all *JK* particles {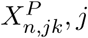 = 1 : *J*, *k* = 1 : *K*} with weights *w*_*n*,*jk*_. This process, resampling two or more layers of the hierarchy at the same time, we call *simultaneous resampling*. The staggered resampling in (S23) can have computational advantages in terms of memory: one never needs to keep all *JK* particles in memory simultaneously. Also, staggered resampling is convenient in a multi-processor computational environment, where the computations for the first layer of the hierarchy can be split across processors and the second layer can be computed without any need for communication between processors.

Another motivation for hierarchical sampling arises when one can separate the generation of the prediction particle into a computationally expensive step followed by a cheap step. Heuristically, if the particles are large and computationally expensive, one wants to ensure that a particle does not get culled due to a single unfortunate draw from a proposal distribution. A component of the proposal distribution that is computationally expensive but not critical for the particle weight should be carried out relatively few times. By contrast, a component of the proposal distribution that is computationally cheap but critical for the particle weight, and hence for the survival of the particle, should be carried out relatively many times. For this motivation, there may be no compelling reason to carry out staggered resampling, in which case simultaneous resampling should be preferred. Both hierarchical sampling possibilities can arise in different parts of a single algorithm, potentially giving rise to several layers of sampling and resampling.

Hierarchical sampling is a standard technique, and theory exists to guide a good sampling structure (Skinner et al., 1989). In practice, however, preliminary experimentation is a good guide. Hierarchical resampling receives diminishing returns for increasing values of *K*, since since *J* is the basic Monte Carlo sample size which asymptotically justifies the Monte Carlo approach. Moderate values of *K* > 1 can have compelling practical advantages, which can be quantified by evaluating the variance of the Monte Carlo likelihood estimate.

### S3.4 Just-in-time evaluation of some state variable components

In equation S3, our GenPOMP model included state processes {Γ(*t*)} and {Δ(*t*)} which have no role in the dynamics, meaning that they do not affect the infinitesimal transition probabilities for {Φ(*t*)} and {Ψ(*t*)} but do affect the measurements. If the measurements depend only on some subset or combination of these state variables, it is computationally desirable to generate the required subsets or combinations only when needed. Carrying out this computational shortcut, which we call justin-time generation, does not change the model under consideration so long as the required variables are properly constructed at the time they become necessary. Two advantages to just-in-time state variable generation are

1. There may be state variables which, on some event of positive probability, have no effect on the measured components of the system. These state variables can be omitted when carrying out inferences on the rest of the system.
2. The sampling of these variables, and consequent resampling of particles, occurs only when information on the just-in-time variables arrives. In combination with hierarchical sampling (Section S3.3), trying multiple copies of the just-in-time variables for each particle can help to prevent particles being lost due to a single unfortunate draw of a random variable.

To formalize the definition of just-in-time variables, we suppose that *X_n_* can be split into two parts, written as

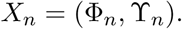

We say that Ξ_*n*_ = *h_n_*(*X*_0:*n*_) gives a just-in-time representation of ϒ_*n*_ if

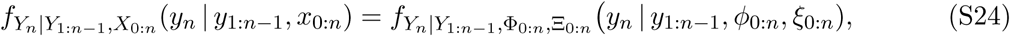

where *ξ_n_* = *h_n_*(*x*_0:*n*_). If we can evaluate (S24) and simulate draws from *f*_Φ*n*,Ξ*n*|Φ_0.*n*–1_, Ξ_*n*–1__, then we can effectively replace ϒ_*n*_ by Ξ_*n*_ in an SMC method such as Algorithm S-2. A particular case, arising in the just-in-time replacement of (Γ(*t*),Δ(*t*) by (*U*(*t*),*V*(*t*)) in Algorithm S-1, occurs when the dynamics of {Φ_*n*_} do not depend on {ϒ_*n*_}, i.e.,

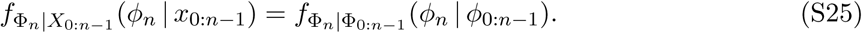

In this case, implementing a just-in-time scheme requires that we can draw from *f*_Ξ_*n*_|Φ_1:*n*_,Ξ_*n–1*__ and we can evaluate the density in equation (S24). In practice, Ξ_0_ may be a trivial random variable, since there is no observation at *t*_0_, but this is not necessary for the just-in-time construction.

The utility of just-in-time evaluation depends in part on the reduction of dimension in replacing Ξ_*n*_ by ϒ_*n*_. For example, nothing is gained by the just-in-time representation Ξ_*n*_ = ϒ_*n*_.

### S3.5 Moving from discrete time to continuous time

Continuous time Markov population models can be approximated in discrete time by a Markov chain (Breto et al., 2009) using a stochastic Euler method. A continuous time measurement model can similarly be discretized to match the time steps of the Euler approximation. For a continuous time latent process model, suppose that {*X*(*t*),*t* ∈ 𝕋} is a right continuous stochastic process taking values in 𝕏. We suppose that the continuous-time measurement process {*Y*(*t*)} consists of a counting process, {*D*(*t*)}, together with a sequence of measurements {*G*_1_, *G*_2_,… } where *G_n_* occurs at time *T_n_* = inf {*t* : *D*(*t*) ≥ *n*}. This notational setup is based on Section S1, but we no not require any of the additional structure of a GenPOMP model at this point. We write 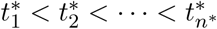 for the observation times of the data, 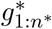. Here, we suppose that *D*(*t*) is part of *X*(*t*) and, specifically, is represented by the observed component *B*(*t*) in the decomposition

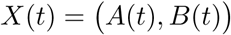

corresponding to a continuous-time version of equation (S18). This situation arises in GenPOMP models when {*D*(*t*)} counts diagnosis events for a disease transmission model {*X*(*t*)}, as in Section S1. Suppose that the rate of observation events at time *t* does not depend on the measurement process {*Y_n_* : *t_n_* ≤ *t*} given the current state process *X*(*t*), i.e.,

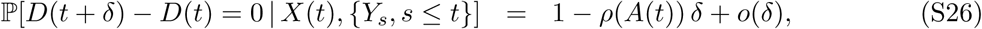

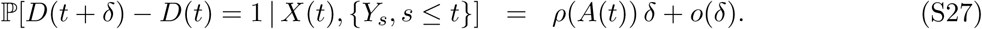

Then, dividing the interval 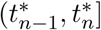 into subintervals of width *δ* and taking *δ* → 0, the limit of discrete approximations using (S26) and (S27) corresponds to a combined weight from evaluating (S23) in each of the 1/*δ* subintervals with no measurement followed by one subinterval with a measurement, i.e.,

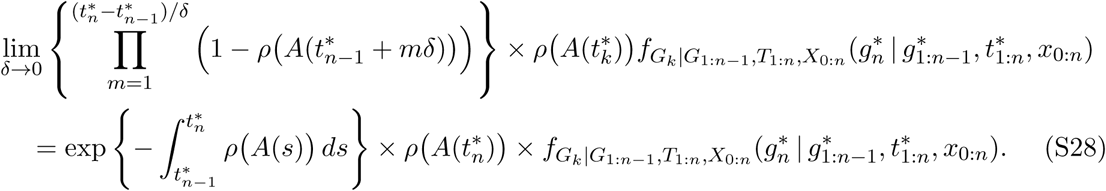

Note that one can view the first two terms of the product in equation (S28) as a density with respect to Poisson counting measure.

## S4 Details of the HIV model used in the main text

In this section we provide additional details that describe the HIV model used in the main text. As the system is Markovian, we can fully specify the model by defining probabilities of each possible change to the state of the system given the current state over an interval of time *δ*. There are three types of events that change the state of system, each in a fundamentally different way:

1. An individual changes class. This event modifies an existing lineage on a transmission tree.
2. An individual in the study population infects a new individual. This event adds a new lineage to an existing transmission tree.
3. An individual outside the study population infects a new individual. This event seeds a new transmission tree consisting of a single individual. The genetic tree associated with with this new transmission tree joins all other genetic trees at the polytomy.

We define probabilities for the first two types of events from an individual-based perspective. Recall that the state of any individual *i* at time *t* is given by a random process {*X_i_*(*t*)}. The probabilities of class changes for each individual over an interval of time *δ* are given by

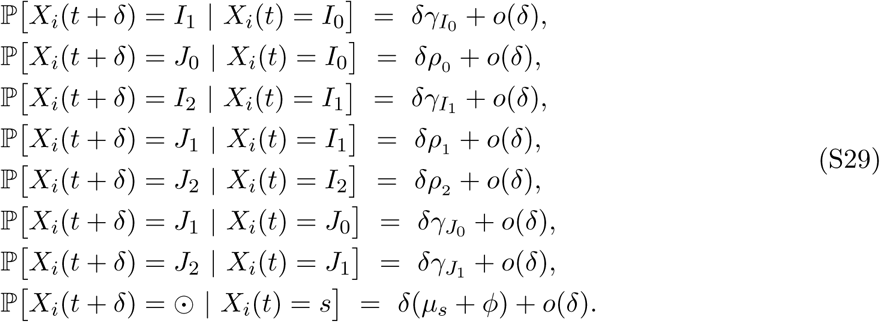

Above, *μ_s_* is a state-dependent death rate for an individual in state *s* ∈ 𝕊 = {*I*_0_,*I*_1_,*I*_2_, *J*_0_, *J*_1_, *J*_2_}, *X_i_*(*t*) = ⊙ if individual *i* is not in the study population at time *t*, and *ϕ* is a constant rate of emigration from the study population. The probability that an infected individual from inside the population gives rise to a new infection is,

𝕡 [the i^*th*^ individual infects a new individual in [*t*, *t* + *δ*]| *X_i_*(*t*) = *s*] = *δ*_*ε*_*s*__ + *o*(*δ*), where *ε_s_* is the infectiousness of an individual in state *s*. The probability that an infected individual from outside the population gives rise to a new infection is, 𝕡 [an infection occurs from outside the study population in [*t*, *t* + *δ*]] = *δϕ* + *o*(*δ*).

Note that this last probability, in contrast to those before, is not defined on a per capita basis. Also note that all new infections start in class *I*_0_; this model does not allow immigration of later stage infected (or diagnosed) individuals into the population.

This model closely resembles a model from a recent phylodynamic analysis of the Detroit HIV epidemic Volz et al. (2013a), but differs in key ways. First, whereas Volz et al. (2013a) modeled incidence as a smooth, deterministic function, we model incidence mechanistically as a function of the states of individuals in the system. Second, instead of using a system of deterministic ordinary differential equations to model counts of individuals in each state, our model incorporates stochasticity into the process of state transitions.

### S4.1 Initial values for the HIV model

The *initial value* for a GenPOMP model is *X*(*t*_0_). In general, the initial value can be treated as an unknown parameter vector which can be estimated using our GenIF methodology. There may be only limited information about these parameters in the data, but that is not a major problem for constructing profile likelihood estimates on other parameters of interest. However, a more parsimonious modeling approach is to set *X*(*t*_0_) to be a suitable function of the values of the dynamic parameters. For example, under a stationarity assumption for the dynamic system, one might set *X*(*t*_0_) to be a random draw from the stationary distribution or some mean value approximation to this. Our HIV model is not stationary, since we follow an age-cohort, but nevertheless we decided to initialize at plausible values given the dynamic parameters rather than estimate additional parameters. Further investigation could relax this assumption.

Part of the specification of *X*(*t*_0_) involves determining the genetic relationship assumed between infections that do not occur in the study population during the modeled period. The time *t*_0_ at which we start modeling the population does not have to match the time at which we start to observe it. We could, for example, have zero sequencing probability before some time point. However, for our HIV model, these two times coincide. In the context of this HIV model, this component of the initial value involves determining the depth of the assumed polytomy, quantified by the time *t*_root_ < *t*_0_ at which all trees in the transmission forest are modeled as meeting in the phylogenetic tree.

We carried out the following construction of the initial values of the membership of each compartment. We first note that the total number of diagnosed individuals is a perfectly observed quantity. By selecting a cohort, we have the advantage of working with a well-defined subpopulation. Over the time period from 2000 to 2012 we know exactly how many individuals were diagnosed. The MDCH dataset only has gene sequences between 2004 and 2012, so we decided to set *t*_0_ = 2004. By 2004, the cohort grew to have 42 diagnosed individuals. Our aim in specifying initial counts is to apportion these 42 individuals to the three different classes of diagnosed individuals and populate the three unobserved states (the undiagnosed individuals) with counts. We assume no deaths over this period of four years. We constructed initial counts for each class by calculating under some additional assumptions under which these values become numerically tractable. First, we made the approximation that all rates of flow, with the exception of *h*(*t*), are fixed at a current parameter estimate. Further, we suppose that *h*(*t*) is constant at some fixed value,

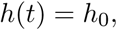

ignoring the dependence of *h*(*t*) on the state of the system. We then approximate the initial state by setting up and solving differential equations representing a deterministic solution to the model equation, formally equivalent to requiring the system of equations (S29) to hold in expectation. We fixed all rates of flow except *h*(*t*_0_) as described in the main text. Then, if the study cohort begins with all counts at zero in 2000, there is only one possible *h*_0_ for which the total number of diagnoses in this approximating model matches the observed total number of diagnoses. We then solve for this value of *h*_0_ and in doing so we obtain the counts in each compartment. Trajectories for the six states and their final values after four years are shown in Figure S4.1. This approach to setting initial counts is not self-consistent with the model, as the model assumes that the rate of new infections is dependent on the state of the system, or with the timing of diagnoses observed in the four years leading up to the start of filtering. This simple way of setting the initial conditions is a starting point. Exploring the effect of initial conditions on model fits could be an area of future work.

We treated the time of the polytomy as an initial value parameter, with each particle starting with its own polytomy time. In this way, the polytomy time fits naturally into the iterated filtering maximization routines.

**Figure S-3:**
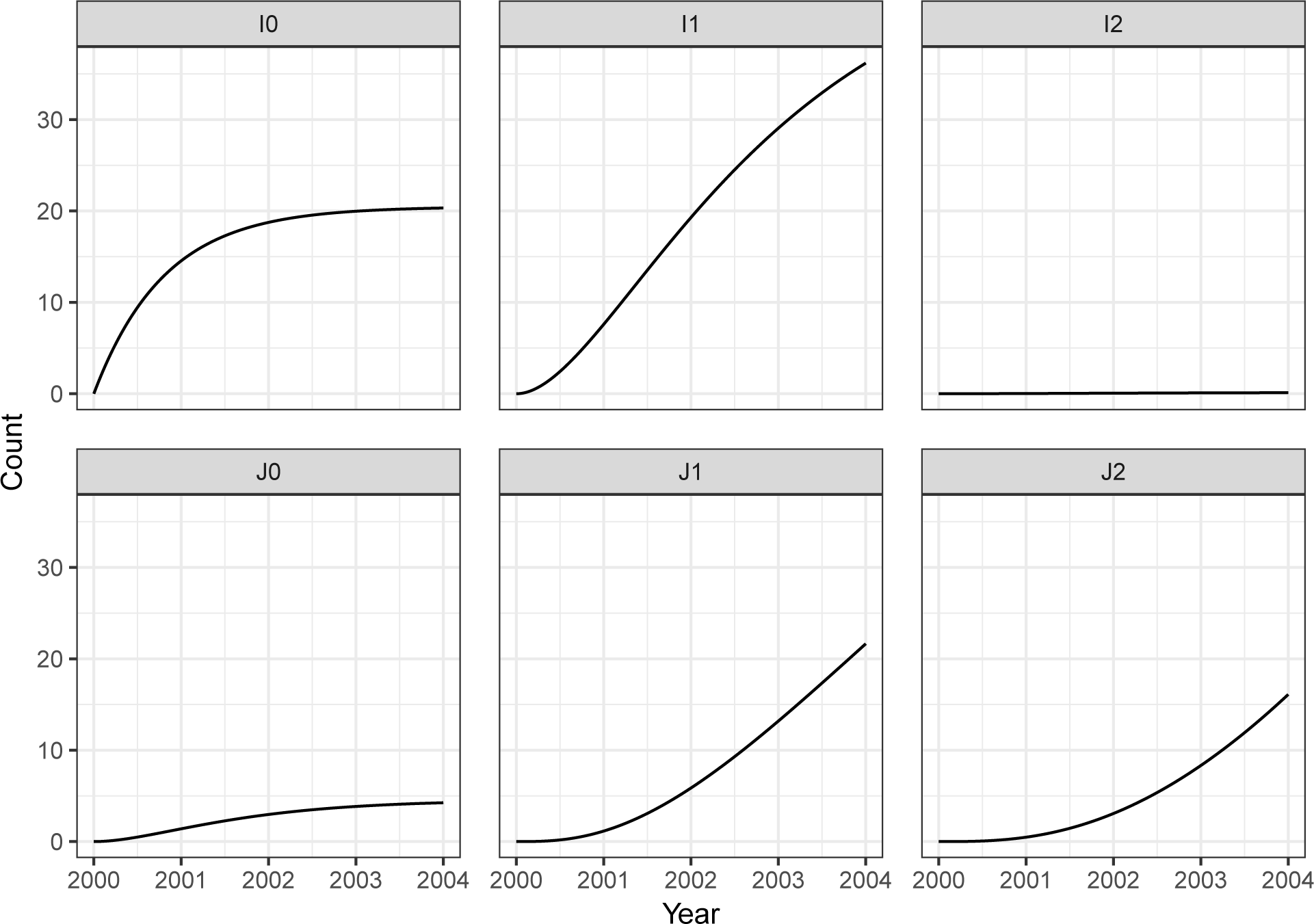
Trajectories of counts of each class of infected individuals over four years prior to *t*_0_ = 2004 when assuming a constant rate of new infections, all flows between and out of compartments as specified in the main text, and zero individuals initially in the cohort. We used the resulting counts in 2004 as the initial values for the data analysis.

### S4.2 Algorithmic parameters used for the numerical results

The choice of algorithmic parameters can affect the numerical efficiency of the GenSMC and GenIF algorithms. For large computations, when Monte Carlo variability is an appreciable component of parameter uncertainty, this can have an effect on the quality of the resulting statistical inferences. In Table S-1 we supply the algorithmic parameters that we used in the simulation study (for GenSMC) and in the data analysis (for both GenIF and GenSMC). We selected *J*, *K*, *L* and *M* such that Monte Carlo uncertainty on parameter estimates and confidence intervals was tolerable (Ionides et al., 2016) and such that runtimes were not prohibitively long.

Three of the algorithmic parameters are only used in GenIF: the random walk standard deviation, *σ_rw_*, the cooling factor, *α_c_*, and the number of GenIF iterations, *I*. Together, these parameters determine the extent to which GenIF shrinks the diameter of the parameter swarm. In the GenIF algorithm, perturbation of parameters over which we are maximizing occurs for each particle just before the proposal step. We perturb the parameters by multiplying each by a random deviate from a log normal distribution with mean one and standard deviation 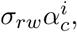 where *i* ∈ {0,1, …,*I* — 1} is the iteration of GenIF. This choice of perturbation is appropriate for nonnegative parameters. Although our framework allows for a different random walk standard deviation for each parameter, in this case we found that the same random walk standard deviation for all parameters was effective, and we report this value in Table S-1.

The algorithmic parameters in Table S-1 together with the source code at https://github.com/kingaa/genpomp are sufficient to reproduce the methodology we apply in our analysis. The HIV sequence data we analyzed are not publicly available, in accordance with our data use agreement with Michigan Department of Community Health.

**Table S-1:** Algorithmic parameters used in the simulation study and the data analysis.

| Algorithmic parameter | Description | Simulation Study |  | Data Analysis |  |  |  |
| --- | --- | --- | --- | --- | --- | --- | --- |
|  |  | GenSMC |  | GenIF |  | GenSMC |  |
|  |  | Diagnosis data only | Diagnosis data and genetic sequences | Diagnosis data only | Diagnosis data and genetic sequences | Diagnosis data only | Diagnosis data and genetic sequences |
| $J$ | Number of particles | 10000 | 60000 | 10000 | 10000 | 10000 | 10000 |
| $K$ | Number of attachment sites per sequence | - | 5 | - | 10 | - | 10 |
| $L$ | Number of relaxed clock gamma samples per attachment site | - | 10 | - | 10 | - | 10 |
| $M$ | Number of relaxed clock beta samples per gamma | - | 1 | - | 1 | - | 1 |
| $\alpha_c$ | Cooling factor | - | - | 0.95 | 0.95 | - | - |
| $\sigma_{rw}$ | Random walk standard deviation | - | - | 0.01 | 0.01 | - | - |
| $I$ | Number of GenIF iterations | - | - | 50 | 30 | - | - |

